# Age-related differences in the cortical motor network during unimanual and bimanual coordination – an EEG study

**DOI:** 10.1101/2025.08.15.670322

**Authors:** P. Babaeeghazvini, B.C.M. van Wijk, J. Gooijers, Cara A. Knott, S.P. Swinnen, A. Daffertshofer

## Abstract

Aging is accompanied by various neurophysiological changes that may affect cognitive and motor function. Compared to younger adults, motor performance in older adults engages more brain regions and exhibits elevated neural activity in the motor-related alpha (8-12 Hz) and beta (15-30 Hz) frequency bands of electro-encephalography (EEG) signals. To what extent these functional changes correlate with the quality of motor performance is underexplored. We recorded EEG in 19 young and 19 older adults during unimanual and bimanual visuomotor tasks with different degrees of difficulty. Older participants showed a lower quality of performance than younger adults, especially during asymmetric bimanual tasks. We analyzed source-localized activity in bilateral parietal and (pre-)motor areas to investigate, especially, the hypothesized pivotal role of left premotor cortex (PMC_L_) activity within the motor network during motor performance. In PMC_L_, beta activity was indeed significantly affected by age during bimanual performance, while alpha activity was altered in the bilateral (pre-)cuneus. When predicting error of performance via alpha and beta modulations, we found significant associations in bilateral M1 and (pre-)cuneus_R_ during unimanual performance, while in bimanual performance, PMC_L_ and (pre-)cuneus_L_ were also included in the significant association. Our results confirm the importance of PMC_L_ and (pre-)cuneus_L_ for the performance of bimanual tasks, especially when the tasks are challenging. The age-related differences in alpha power in bilateral (pre-)cuneus and their associations with motor performance suggest that altered visuomotor integration has an important contribution to the reduction of the quality of motor performance in older adults.

## Introduction

Aging is often accompanied by increasing difficulties in performing coordinated bimanual and multi-joint movements (<u>Seidler et al., 2010</u>). This may jeopardize activities in everyday life like tying shoelaces, eating with a knife and fork, and many other coordinated motor activities (<u>Krehbiel et al., 2017</u>; <u>Swinnen, 1998</u>). An age-related decline in upper limb motor performance may have various origins, from muscle atrophy to changes in the nervous system at different levels. For example, voxel-based morphometry has revealed changes in gray and white matter macrostructure, and changes in white matter tracts (<u>Ding et al., 2016</u>), including myelination (<u>Benitez et al., 2018</u>; <u>Faizy et al., 2020</u>; <u>Kiely et al., 2022</u>), structural connectivity, and their microstructural organization as identified using dMRI (<u>Coelho et al., 2021</u>; <u>Fujiyama</u> <u>et al., 2016b</u>). Additionally, neurotransmitter concentrations may vary in older adults (<u>Burianová et al., 2020</u>; <u>Richardson et al., 2021</u>), potentially affecting synaptic activity (<u>Lall et</u> <u>al., 2021</u>). While all these changes are typically associated with altered brain function and muscular activity (<u>Larsson et al., 2019</u>), it remains unclear how they collectively influence the control of coordinated motor activity (<u>Bangert et al., 2010</u>; <u>Maes et al., 2017</u>; <u>Swinnen, 1998</u>).

Intra- and inter-hemispheric interactions between (bilateral) pre-motor cortex (PMC) and primary motor cortex (M1) are crucial when performing uni- or bimanual coordination becomes more challenging (<u>Chettouf et al., 2020</u>), and especially the PMCs appear to be more strongly activated in older adults during bimanual coordination performance compared to younger adults (<u>Goble et al., 2010</u>; <u>Heitger et al., 2013</u>; <u>Larivière et al., 2019</u>; <u>Ward, 2006</u>). Dual-site TMS revealed that older adults are seemingly less successful in regulating the interaction between the left dorsolateral prefrontal cortex and right M1 during bimanual performance, particularly during non-isofrequency conditions, such as when the left hand moves three times faster than the right hand (<u>Fujiyama et al., 2016b</u>). By the same token, both inhibitory and facilitatory inputs from left dorsal premotor cortex (PMd) to left M1 are decreased in older adults compared to younger adults (<u>Ni et al., 2015</u>). To compensate for the altered connectivity between PMd and M1, older adults may show stronger BOLD activity in PMd compared to younger adults (<u>Ni et al., 2015</u>; <u>Wu and Hallett, 2005</u>). While the loss of connectivity between PMd and M1 might underlie fundamental motor functions like planning, preparation, and execution (<u>Hoshi and Tanji, 2007</u>) as well as coordination (<u>Ward, 2006</u>), the extent to which a reduced functionality of (especially the left) PMC is associated with the quality of motor performance in older adults is not well understood. In electro-encephalography (EEG) assessments (here, our primary approach), one can expect to find reduced motor task-related alpha (8-12 Hz) and beta (15-30 Hz) power in older adults (<u>Rueda-Delgado et al., 2019</u>), which would align with stronger (BOLD) activity during motor execution (<u>Formaggio et al., 2008</u>), especially in premotor areas (<u>Chettouf et al., 2022</u>).

Daily life activities often rely on the integration between motor commands and visual information. Motor performance that includes visual feedback (especially visual tracking) is accompanied by activity in occipital areas and (pre-)cuneus (<u>Rossit et al., 2013</u>; <u>Thut et al., 2006</u>). Like the activity of the motor network, the visual activation patterns are also affected by aging. For instance, <u>Godde et al. (2018)</u> found significantly more activation, i.e., elevated blood oxygen level dependent (BOLD) responses, in the cuneus for older adults compared to younger adults during the exertion of isometric force in precision grip tasks with the right hand to track a target on the screen that moved along a sine wave. On the other hand, an earlier study by <u>Aizenstein et al. (2004)</u> reported more widespread, negative BOLD responses in cuneus and precuneus when older adults conducted a simple bimanual visuomotor task (pressing a key with both index fingers in response to the visual presentation of a word). However, these widespread negative BOLD responses in older adults were interpreted as the result of more unconstrained visual processing during rest in the scanner. Considering that stronger BOLD responses often correlate with reduced power in EEG (<u>Formaggio et al., 2008</u>; <u>Hanslmayr et al.,</u> <u>2011</u>; <u>Yuan et al., 2010</u>), we hypothesized that age-related decreases in alpha and beta power in the parieto-occipital regions are associated with visuomotor performance quality.

Our aim was to assess age-related differences in both alpha and beta power modulations in motor and visual cortex during unimanual and bimanual visuomotor tasks, and their associations with the quality of motor performance. EEG was used to detect subtle alterations of cortical activity at high temporal resolution, allowing for the discrimination between event-related synchronization (ERS) and desynchronization (ERD) in different frequency bands. Previous studies in young, healthy adults already demonstrated alpha/beta ERD during dynamic force production, while ERS occurs during static force production (<u>Nakayashiki et al., 2014</u>). Interestingly, alpha and beta ERD have already been shown to increase with age in parietooccipital and motor regions (<u>Heinrichs-Graham and Wilson, 2016</u>; <u>Inamoto et al., 2023</u>), while ERS seems to be reduced (<u>Hübner et al., 2018</u>; <u>Inamoto et al., 2023</u>; <u>Liu et al., 2017</u>). To investigate how these age-related spectral power changes are associated with the quality of motor performance, we adapted an experimental design introduced by <u>de Vries et al. (2016)</u> in which participants generated isometric forces using a precision grip to track a force target displayed on a computer screen. Since our experimental tasks involved dynamic force production followed by static force production in a timely manner, a rapid transition from ERD to ERS was deemed necessary to achieve accurate performance. We expected to find lower overall (mean) alpha and beta power during task performance as an indication of stronger cortical involvement in older adults relative to younger ones, as well as possibly a reduced power modulation (i.e., differences between ERD and ERS power) as an indication of a reduced capacity to switch between motor states. We hypothesized that these effects would be most prominent in the left PMC (<u>Babaeeghazvini et al., 2018</u>; <u>Heinrichs-Graham and Wilson, 2016</u>; <u>Xifra-</u> <u>Porxas et al., 2019</u>). We also hypothesized the age-related reduction in mean power and power modulation in the left PMC to be negatively associated with the quality of motor performance.

## Methods

### Participants

Two groups of healthy adults participated in this study (19 younger adults: 9 females, age range 20-30 years, mean=25.7, SD=3.5; and 19 older adults: 10 females, age range 58-75 years, mean=66.4, SD=5.3). Except for two young and one older adult, all were self-reportedly right-handed. None of them had professional musical training or any known neurological, psychiatric, or chronic somatic disease history. The local ethics committee of the faculty of Behavioral and Movement Sciences, Vrije Universiteit Amsterdam, approved the study (2013-45M). All participants provided written informed consent before partaking.

### Protocol

Participants were comfortably seated in front of a computer screen (diagonal 21", resolution 3440 × 1440 pixels, frame rate 100 Hz) with their forearms resting on a table (Figure 1). They were asked to squeeze the tips of two compliant force sensors with the left- and/or right thumb and index finger. The amount of generated force was displayed in real-time on the computer screen with a cursor (blue dot, 15 mm diameter). The goal was to follow a target force (red dot, 35 mm diameter) as accurately as possible in space and time by applying force to the force sensors to control the movement of the cursor. Each trial lasted 8 s, and there was a 2s rest (inter-trial interval) between consecutive trials. The 2s rest was followed by a go-cue after which the target starting to move continuously for 5s with a speed of 100 pixels/s to the final target position in the middle of the screen (ramped force; referred to as *dynamic force*), where it stayed fixed for the remaining 3s (constant force; referred to as *static force*) and followed by another 2s rest. The motor task, hence, lasted a total of 8s. Across tasks and conditions, maintaining the final position implied generating a force of 1.8N on each force sensor (see also Figure 2 below). 64-channel EEG and surface EMG from the first dorsal interosseous and flexor pollicis brevis muscles were recorded simultaneously throughout the experiment.

**Figure 1.**
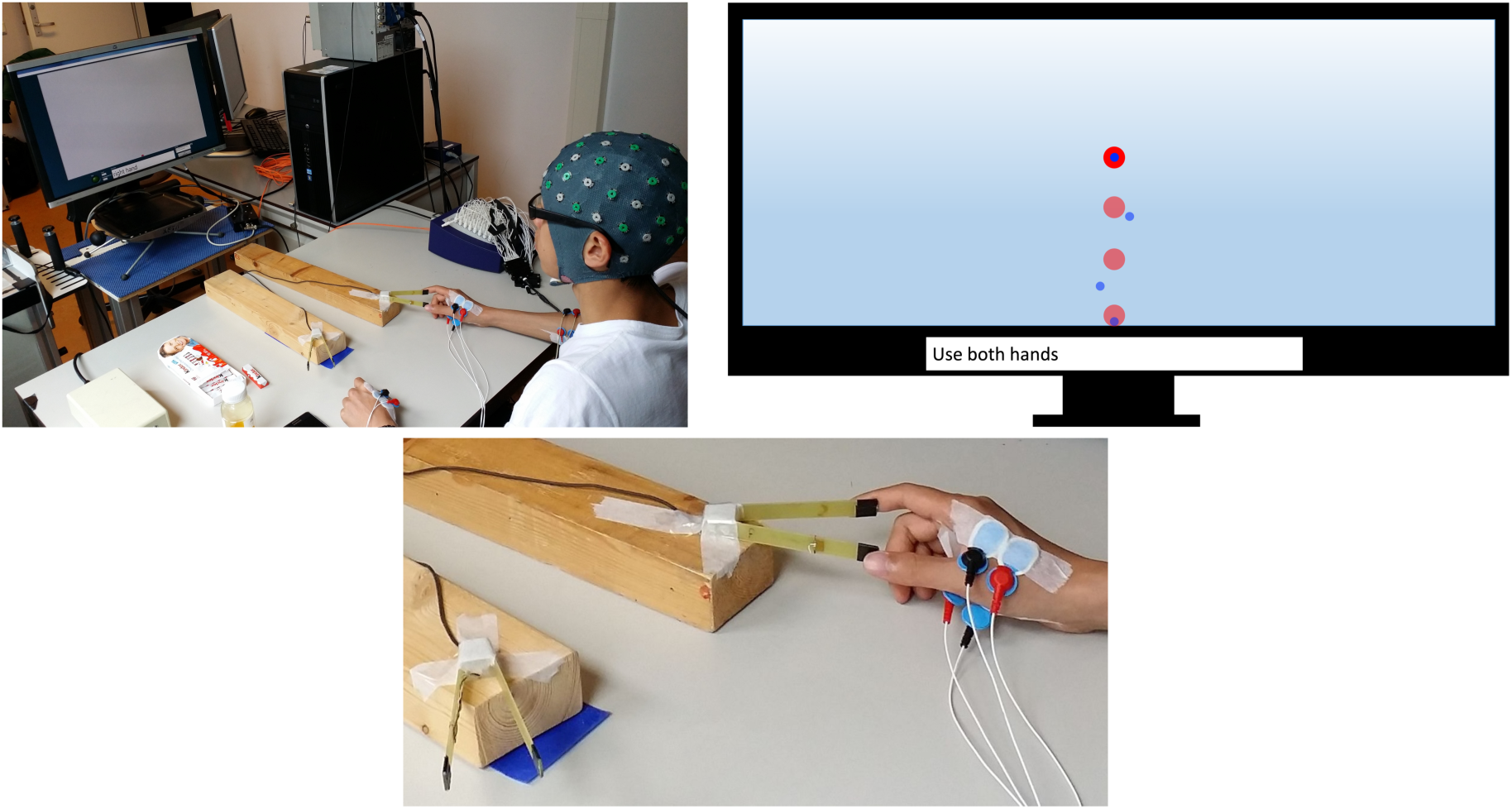
Participants applied dynamic and static pinch grip forces with their index fingers and thumbs to position a cursor on a computer screen. Forces were collected using a custom-made force sensor (bottom panel). During the task, a target (top right panel, red dot) moved upwards continuously on a straight line for 5s with a fixed speed, and stopped just above the middle of the screen for 3s. Participants were instructed to move the cursor (top right panel, blue dot; by applying appropriate force to the sensors) as closely as possible to match the target. The top left and bottom panels show the task set-up and sensors.

**Figure 2.**
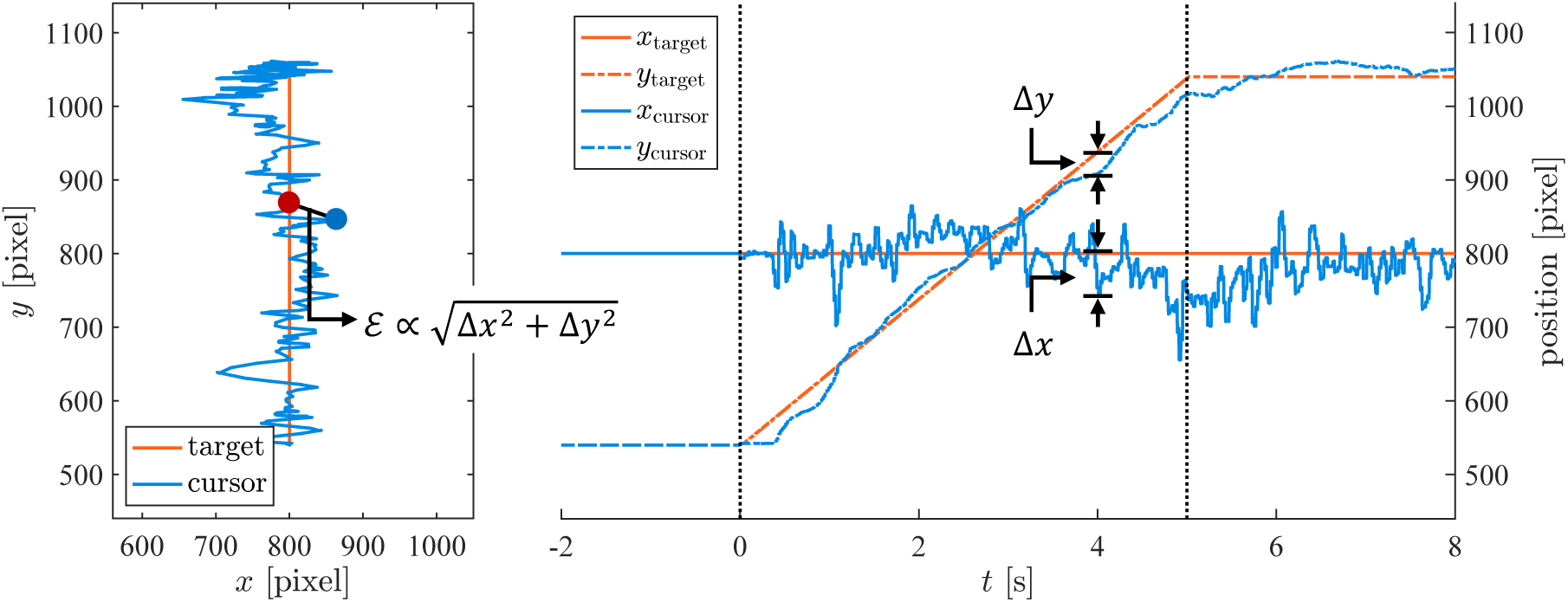
The left panel illustrates the trajectories of the target (orange) and an exemplar trajectory of the cursor (blue); cf. Figure 1, top-right panel. We measured performance via the Euclidean distance between target and cursor. For this we determined the corresponding differences Δ*x* and Δ*y* along the horizontal and vertical axes at every time point (right panel). For a similar illustration of the left/right force errors (*Δf*_*t*_ and *Δf*_*R*_), see also Figure A.1 in the *Appendix*.

Participants conducted unimanual and bimanual tasks under different conditions and at different difficulty levels. During (normal or difficult) unimanual tasks, pressing the corresponding (left or right) sensors moved the cursor straight upwards (90°) (<u>de Vries et al., 2016</u>). During bimanual tasks, pressing the right (left) force sensor moved the cursor to the right (left) upper corner by 45° or 30° under normal or difficult conditions, respectively. In difficult conditions, in addition to 30° of movement, we also artificially increased the sensitivity of one or both sensors by increasing the effect of one or both hand tremor (high frequency) on the screen, as explained in detail below. The experiment comprised a total of eight task/condition combinations (Table 1): four unimanual tasks (two left hand only and two right hand only tasks, with one under the normal and one under the difficult condition for each hand) as well as four bimanual tasks: one with both hands under normal conditions and one with both hands under the difficult condition (both symmetric), as well as one with the left hand under the normal and the right hand under the difficult condition, and vice versa (both asymmetric).

**Table 1.**
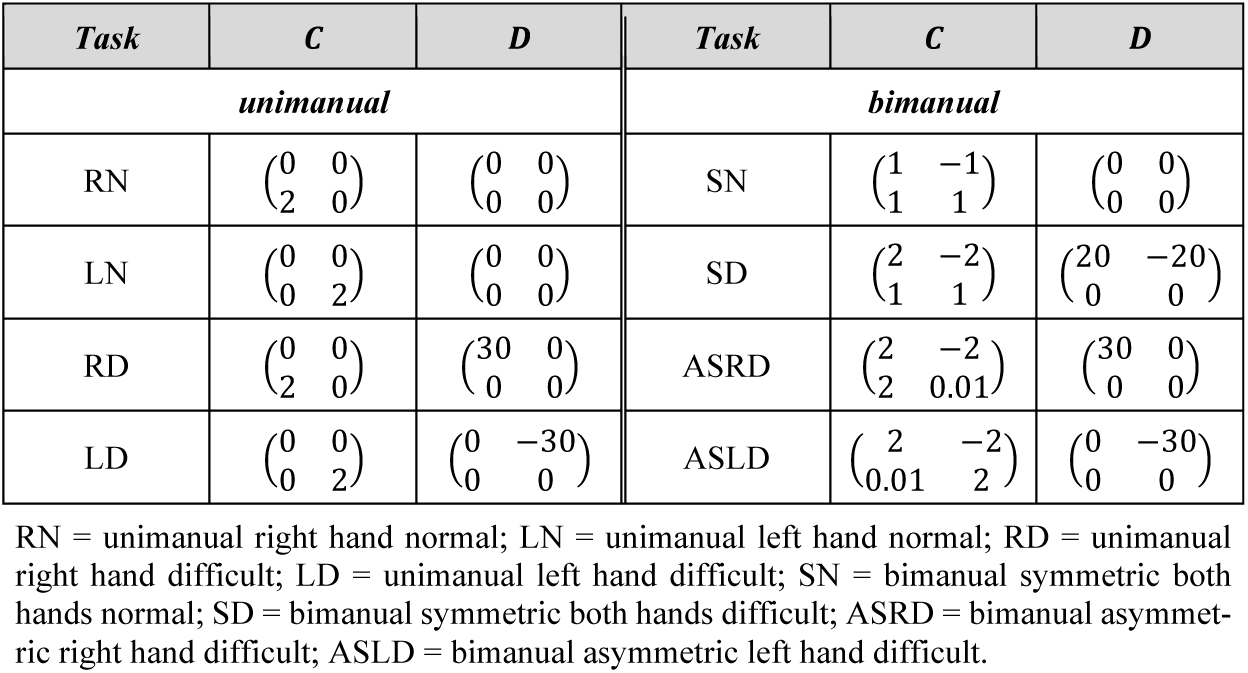
Definition of the different tasks and force mappings to the screen using the mixture matrix *C*, with and without tremor amplification via the matrix *D*.

An experimental session consisted of three recording sets, with several blocks that contained 10 trials of one of the eight conditions. There were 20 blocks in total, with 3 blocks for each of the 4 bimanual conditions and 2 blocks for each of the 4 unimanual conditions. These 20 blocks were placed in 3 sets in a way that the first two sets included seven blocks, and the last set contained six blocks. In a set, blocks were presented in random order. After each block, a 7s rest period was included. Between each set, a short break of several minutes was included to let the participants relax. Task instructions with regards to which hand(s) to use were presented at the bottom of the screen. This information was presented from 3s prior to the start of the first trial until the end of the block. Prior to the main experiment, participants practiced randomly selected tasks for familiarization, including two blocks of unimanual performance (normal and difficult conditions; with each hand performing one of the practice blocks), one block of symmetric bimanual performance (both hands under either normal or difficult conditions), and one block of asymmetric bimanual performance (either bimanual asymmetric right hand difficult or bimanual asymmetric left hand difficult). Participants could repeat this phase until they felt comfortable performing the task.

#### Task difficulty

Participants had to position the cursor on the screen in line with the target. When summarizing the corresponding screen coordinates as a column vector, the target position can be cast in the form of 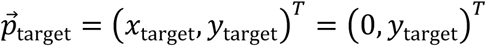 where we note that *x*_target_ was always set to zero, i.e., the target moved upwards without horizontal deviation (superscript *T* denotes the transpose). The cursor’s position was defined as a function of the generated forces 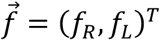 given by

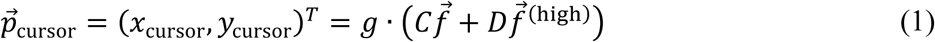

where subscripts *R* and *L* imply the right and left hand, respectively. In other words, the forces 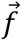 were linearly mapped to the screen using a mixing matrix *C*. The mapping was extended with additional mixture matrix *D* to increase task difficulty by adding the amplified high-frequency components 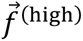 (above 3 Hz) of the measured force signals to the cursor to amplify the high frequency movements. This was meant to artificially increase the effect of hand tremor on the screen and hence to demand more precise force application via pinch grip. We chose *D* to act along the horizontal axis. The scalar gain factor *g* was 90 pixels/N. While all the task-specific mixture matrices are summarized in Table 1, we briefly illustrate a few examples.

In the unimanual right hand normal (RN) task without tremor amplification, *C*_21_was equal to 2, and all other elements were set to 0, implying 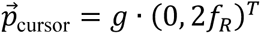. In the unimanual left hand normal (LN) task *C*_22_was equal to 2. In the bimanual symmetric hands normal and difficult tasks (SN & SD), the matrix elements *C*_*i*j_ were set in such a way that they required identical force levels from each hand to produce the vertical displacement. In the absence of tremor amplification, we used *C*_11_ = −*C*_12_ = 1 and *C*_21_ = *C*_22_ = 1 in the SN task, i.e., 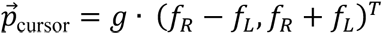. The tremor amplification only affected the horizontal positioning of the cursor (hence *D*_21_ = *D*_22_ = 0), and it was only used in the unimanual right-hand difficult (RD), unimanual left-hand difficult (LD), bimanual symmetric both hands difficult (SD), bimanual asymmetric right-hand difficult (ASRD), and bimanual asymmetric left-hand difficult (ASLD) tasks. For example, in the RD task, we used *D*_11_ = 30, *D*_12_ = 0, to increase displacement in the vertical direction, but the total required force remained largely identical to the case without tremor amplification because in both conditions *C*_21_ = 2. In this case, we used 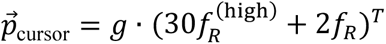. In the SD task, the tremor of both hands was amplified by setting *D*_11_ = 20 = −*D*_12_; see Table 1.

As said, adding 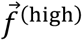 effectively amplifies the high-frequency tremor, which is expected to deteriorate motor performance. In the bimanual tasks, we further hypothesized that due to interhemispheric crosstalk in the ASRD and ASLD tasks, i.e. when 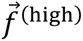 is added only for one side, we will observe a reduced quality of performance at the side where the high-frequency tremor is not artificially amplified.

### Data acquisition

#### Behavioral data

Pinch-grip forces were collected through custom-made force sensors with a strain gauge (type: FAE2-A6174J-35-S13E), allowing for application of up to 3.5N. A bridge amplifier (NI SCXI-1121; ADD National Instruments) served for data acquisition and provided task synchronization triggers. Force data were sampled at a rate of 1,000 Hz. Online high-pass filtering (2^nd^-order Butterworth) was applied to amplify the force signals above 3 Hz whenever appropriate (see above).

#### Cortical activity

EEG was recorded using a 64-channel amplifier (Refa, TMSi, Enschede, the Netherlands; Ag/AgC1 electrodes mounted in an elastic cap according to the extended 10-20 system) with average reference. Signals were digitized with a 2,048 Hz sampling rate. The conductance gel kept the contact impedance of the electrode-skin interface below 10 kΩ.

We would like to note that we also recorded bipolar electromyography of selected muscles (see Figure 1), but we will not discuss this further in view of our focus on the relation between cortical activity and motor behavior.

### Data Analysis

#### Behavioral data

To assess the error of performance (ℰ), we determined the Euclidean distance between the position of the target and the cursor on the screen as illustrated in Figure 2.

The corresponding mathematical form can be given as

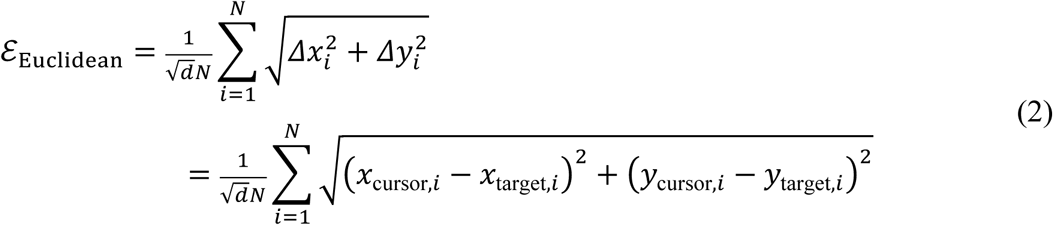

where *d* denotes the degrees of freedom, i.e., *d* = 1 for unimanual and *d* = 2 for bimanual tasks, and *N* denotes the number of samples in time. We also estimated the left force and right force error compared to the target force to identify possible effects on the performance of one hand when manipulating the degree of difficulty of the other during the bimanual tasks.

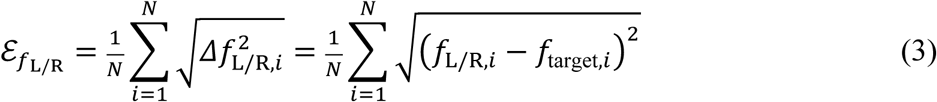

In both cases, the ℰ-values were averaged over trials per participant for each condition and then log-transformed before entering statistical assessments.

#### Electrophysiological data

Data (pre-)processing and analyses were performed in MATLAB (version 2022b, The MathWorks, Natick, MA), including the open-source FieldTrip toolbox (<u>Oostenveld et al., 2011</u>). The behavioral signals were up-sampled to 2,048 Hz via linear interpolation to agree with the EEG and temporally aligned using co-registered TTL-pulses. Possible sampling drifts were corrected using linear interpolation, where the EEG sampling served as a reference timeline.

#### Preprocessing

Data cleaning involved multiple steps. (1) Channels were considered bad when, across the entire recording, they displayed flat lines or when the signal’s mean or standard deviation exceeded 500 mV or 20 times the median standard deviation over all channels. Bad channels were interpolated using the signals from the neighboring channels (linear interpolation and neighbors defined through triangularization of the EEG cap). (2) Per channel, possible signal jumps were defined as an increment between two samples that exceeded 3 times the signal’s standard deviation over all channels after subtracting the 2s median filtered signal. The corresponding time points ±1 ms were linearly detrended and jump points were corrected using shape-preserving splines. (3) Per channel, outliers (spikes) were defined as samples that exceeded 10 times the signal’s standard deviation for that channel or 500 μV after (a) high-pass filtering (10 Hz, 2^nd^-order bidirectional Butterworth) or (b) after subtracting the 3s median filtered signals (both conducted separately followed by independent spike detection); the corresponding ±2 ms samples were corrected via shape-preserving spline interpolation of the unfiltered data. (4) Power line artifacts were removed via notch filtering (3^rd^ order bi-directional Butterworth filter, band half-width 0.1 Hz) at 50 Hz and its harmonics up to the Nyquist frequency (1,024 Hz). (5) Data were decomposed using independent component analysis (ICA). Independent components were considered an artifact if they were dominated by the channels FP1, FP2, FPz, O1, O2, and Oz (eye movements), if their median frequency was below 5 Hz (head movements or eye blinks), or if their median frequency exceeded 100 Hz (muscle/EMG artifacts, based on the expected median frequency of, e.g., the masseter muscle). On average, we removed 4.8 components due to eye movement, 0.3 due to head movement and eye blinks, and 2.7 due to muscle activity per participant (see the *Supplementary Material* for more details related to the number of removed ICA components per participant in Table S.1). (6) All channels were finally checked for residual spikes exceeding 250 μV. If present, the corresponding channels were epoched according to the experimental design from 1 to 8s after the event trigger at *t* = 0s, which indicates the start of a new trial or event (*t* = 0s refers to the moment of the gocue). Epochs with peaks > 200 times the standard deviation over time of the event-related average were considered bad, and the corresponding events were removed. After cleaning, the EEG signals were band-pass filtered between 3 and 250 Hz using a 3^rd^-order bi-directional Butterworth filter and re-referenced to the average over channels.

All pre-processed signals were down-sampled to 512 Hz after correcting for aliasing using a bidirectional FIT filter at the Nyquist frequency (256 Hz). The signals were finally segmented into separate trials of 9s each, with the first 1s of pre-movement (planning for movement), 5s of dynamic target tracking (dynamic force), and 3s of static grip force (static force); we note that the time point 0s always corresponds to the onset of dynamic target tracking.

#### Sensor-level analysis

To first identify the relevant epochs and frequency bands, we conducted a time-frequency analysis over occipital/parietal and fronto-central channels using conventional Morlet wavelets (4-80 Hz, in steps of 0.25 Hz, width = seven cycles, length = three standard deviations of the implicit Gaussian kernel and time resolution of 2/(sampling rate)). Spectra were normalized by dividing per frequency by the average power (the mean standard deviation of power over all conditions, channels, and time points) within the [1-8]s interval (we chose the epoch of 1 to 8 seconds because several participants started performing with some delay). Pooling the wavelet spectra of all participants and conditions (of both young and older adult groups) revealed that the differences between the static and the dynamic force were particularly pronounced (with static power higher than dynamic power) in the beta band (15-30 Hz) when looking at channels of the motor strip (fronto-central) and in the alpha band (8-12 Hz) when looking at occipital/parietal channels; see Figure 3.

**Figure 3.**
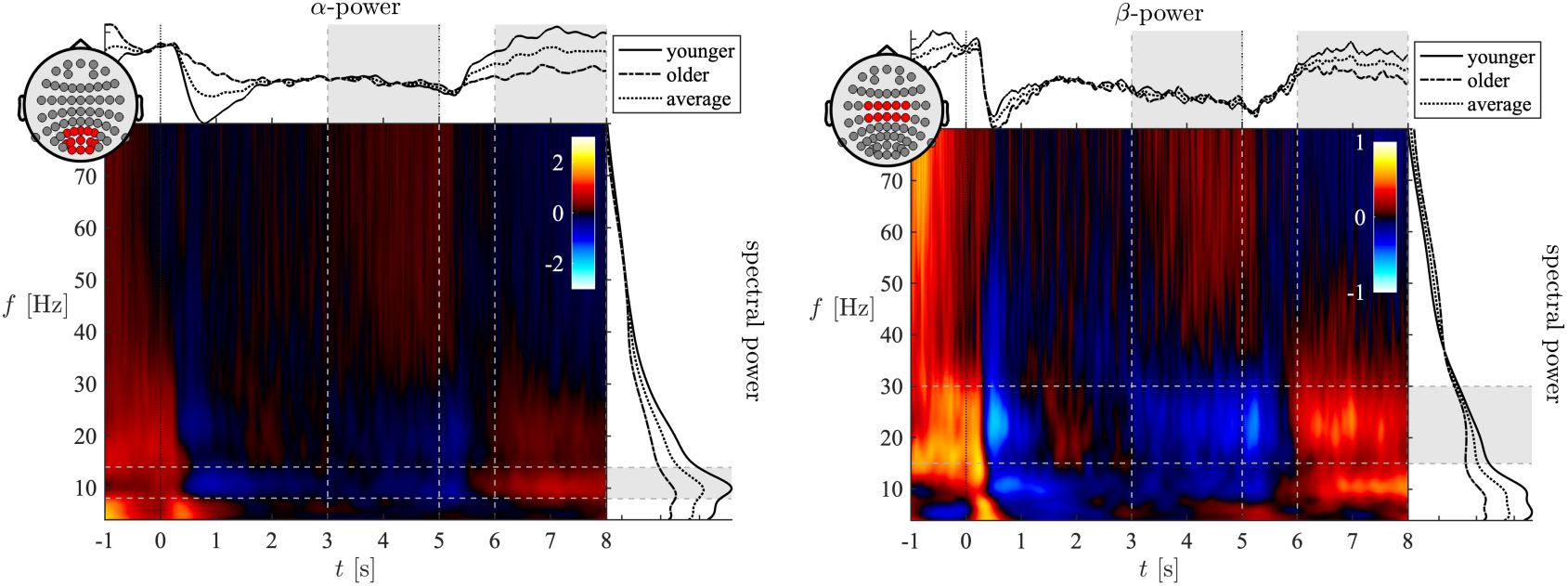
Wavelet spectra of occipital/parietal and frontocentral channels (left panel: O1, Oz, O2, PO5, PO3, PO1, POz, PO2, PO4, PO6, P3, P1, Pz, P2, and P4; right panel: C3, C1, Cz, C2, C4, FC3, FC1, FCz, FC2, and FC4;). We computed the average over trials per condition, averaged all conditions per participant, and averaged over all participants within a group. The time-frequency spectra are the averages over all participants (normalized per frequency to the [1-8]s interval), whereas the log-power spectra (right side of the TF figures; 2D lines) and time-dependent alpha-/beta-power (on the top of the TF figures; 2D lines) are group specific; here they are normalized to the [3-5]s interval to ease visual inspection. The gray bars indicate the epochs and frequency bands of interest. As it is evident from the top 2D lines, older adults displayed lower alpha and beta power in the static [6-8]s phase and the overall power (2D lines on the right side of TF) was also reduced in both frequency bands.

This sensor-level analysis served to identify the exact time and frequency windows for source localization analyses. In consequence, we divided each trial into two epochs: [3-5]s for the dynamic force production phase and [6-8]s for the static force production phase, respectively. As illustrated in Figure 3, the power in the dynamic phase of force production showed a marked reduction in the alpha and beta frequency bands (from 8 to 30 Hz) compared to the static phase. We would like to note that by this choice of time windows for dynamic and static epochs, we did not only avoid any transient behavior (i.e., initiating force production movement too late and/or terminating the force production movement too early) but also ensured identical epoch duration (2s for both the dynamic and static force production phase), reducing possible estimation biases in any of the subsequent statistics.

#### Source estimation

A source model was obtained from the ICBM152 template MRI provided by the FieldTrip toolbox, using the Simbio implementation in FieldTrip (<u>Geweke, 1982</u>; <u>Vorwerk et al., 2018</u>). In brief, a 5-tissue head model was created. We assigned conductivity values of 0.33, 0.14, 1.79, 0.01, and 0.43 S/m to grey matter, white matter, cerebrospinal fluid, skull, and scalp, respectively (<u>Buchner et al., 1997</u>), and estimated a lead field for every grid point (2 mm resolution) via finite elements. All coordinates will be reported in Montreal Neurological Institute (MNI) standard space. Then, using dynamic imaging of coherent sources (DICS) beamformers (<u>Gross et al., 2001</u>), source estimation was performed in the alpha (8-14 Hz) and beta bands (15-30 Hz) per participant across conditions. Moreover, we found the most pronounced differences between dynamic and static forces in the spectral power of these two frequency bands relative to the other frequency bands (see Figure 3). To identify the corresponding spatial filters, we estimated the cross-spectral density *S*_*xy*_(*f*) via discrete prolate spheroidal sequence tapers (DPSS) at the central frequency *f* of the band of interest (we used ± half the bandwidth for spectral smoothing) over the epoch [1-8]s across conditions. Subsequently, we estimated the power in the separate epochs of 3-5s and 6-8s at every voxel *k* in the brain, yielding 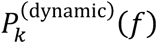 and 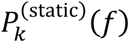. From these, we computed a proxy for the power modulation in terms of a pseudo-*t*-value that we defined as

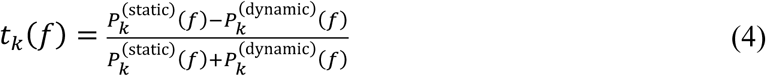

This value quantifies the (normalized) difference between the static and dynamic phases, or between ERS and ERD, respectively (<u>Nakayashiki et al., 2014</u>). To stabilize variance, we applied Fisher’s z-transform to *t*_*k*_(*f*) prior to statistical assessments.

The assessment of the *t*_*k*_(*f*)-values revealed significant voxels (within and between subject statistics) that let us to define six regions-of-interest (ROIs) that we assigned to anatomical regions via the Harvard/Oxford cortical atlas (<u>Desikan et al., 2006</u>); cf. Figure 4: bilateral precentral gyrus, superior frontal gyrus + supplementary motor area, precuneus + cuneus, or in short left/right M1, PMC and (pre-)cuneus, respectively.

**Figure 4.**
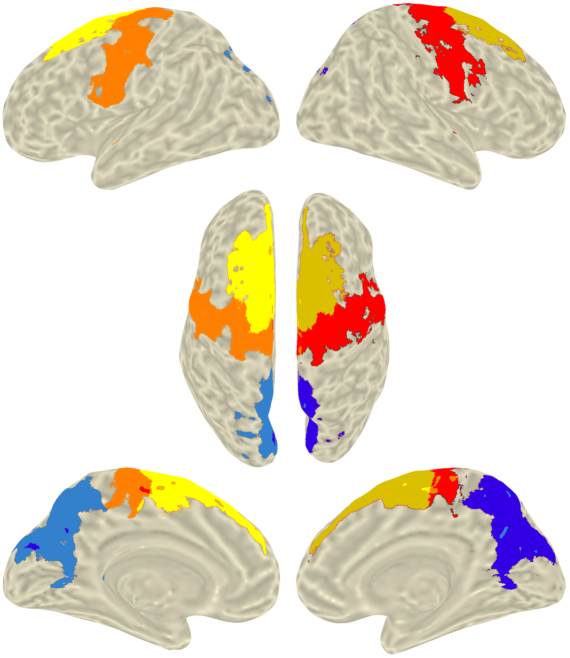
Illustration of the ROIs. The colored areas are the M1 (orange and red), PMC (bright and gold yellow) and precuneus/cuneal cortex (blue and navy). In our analysis the ROIs have been identified as task-related clusters of statistically significant voxels within the corresponding parcellation of the Harvard/Oxford cortical atlas. Mean power and pseudo-*t*-value were estimated for the mean virtual signal over significant voxels within each ROI. The ROIs are overlaid on the pial surface of the MNI 152 template. See Figure 5 for the beamformer results.

Within the motor regions, we selected significant voxels related to the within-subject statistics (effect of task) in the beta frequency band, whereas for the occipital areas we selected significant voxels related to between-subject statistics (effect of age) in the alpha frequency band (see below). Then, we estimated the corresponding source signals using spatial filters optimized over the [1-8]s epoch in the 4-80 Hz frequency band. The optimal dipole orientation per voxel was estimated using principal component analysis (<u>Westner et al., 2022</u>). Per ROI, we chose the mean spatial filter over significant voxels with positive *t*-values within an ROI after weighting them with the respective *t*-value from the contrast between task and resting state (in beta frequency) or between groups (in alpha frequency)– see the *Statistics* section below.

For the source signals in each of the six ROIs, we determined the normalized spectral power using Morlet wavelets (with the same normalization approach that we used for sensor level) to indicate the validity of our source reconstruction.

#### Outcome parameters

For statistical analysis, we defined two outcome parameters per ROI and frequency band: The mean power over the static and dynamic phases is given by

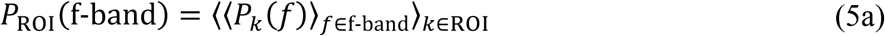

with 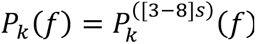, and the mean pseudo-*t*-value is given by

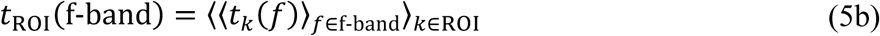

with *t*_*k*_(*f*) given in Eq. (4) and *k* referring to the ROI. The pseudo-*t*-value was meant to target the expected age-related difference in ERD/ERS differences (<u>Heinrichs-Graham and Wilson,</u> <u>2016</u>; <u>Inamoto et al., 2023</u>; <u>Xifra-Porxas et al., 2019</u>).

The power values *P*_*k*_(*f*) were log-transformed, and the pseudo-*t*-values *t*_*k*_(*f*) were Fisher z-transformed prior to averaging over each ROI and over trials per participant and condition.

### Statistics

#### Source estimation

To determine ROIs in the beta frequency band, we tested which voxels reached statistical significance based on a within-subject comparison estimating the effect of tasks against resting state over the entire brain. From the significant voxels related to task performance, we used the mean pseudo-*t*-values over all tasks and contrasted them against zero (considered as baseline; cf. Eq. (4) for the definition of the pseudo-*t*-values). Then, we employed a Monte Carlo approach with a cluster-based permutation statistics where the corresponding distribution was built using 2^14^ permutations. We used threshold of α_cluster_ = 10^-7^ to identify significant clusters, represented by sum of their compound values (<u>Nichols and</u> <u>Holmes, 2002</u>). This was followed by an independent *t*-test (significance threshold α = 5·10^-4^) to identify sources across all participants that, as expected, were predominately located in motor areas (see Fig. 5). In the alpha frequency band, we contrasted younger adults against older adults to estimate the *age* effect on pseudo-*t*-values by using between-subject statistics and estimating the significant voxels resulting from the pseudo-*t*-values with α_cluster_ = 10^-3^ and α = 5·10^-2^.

**Figure 5.**
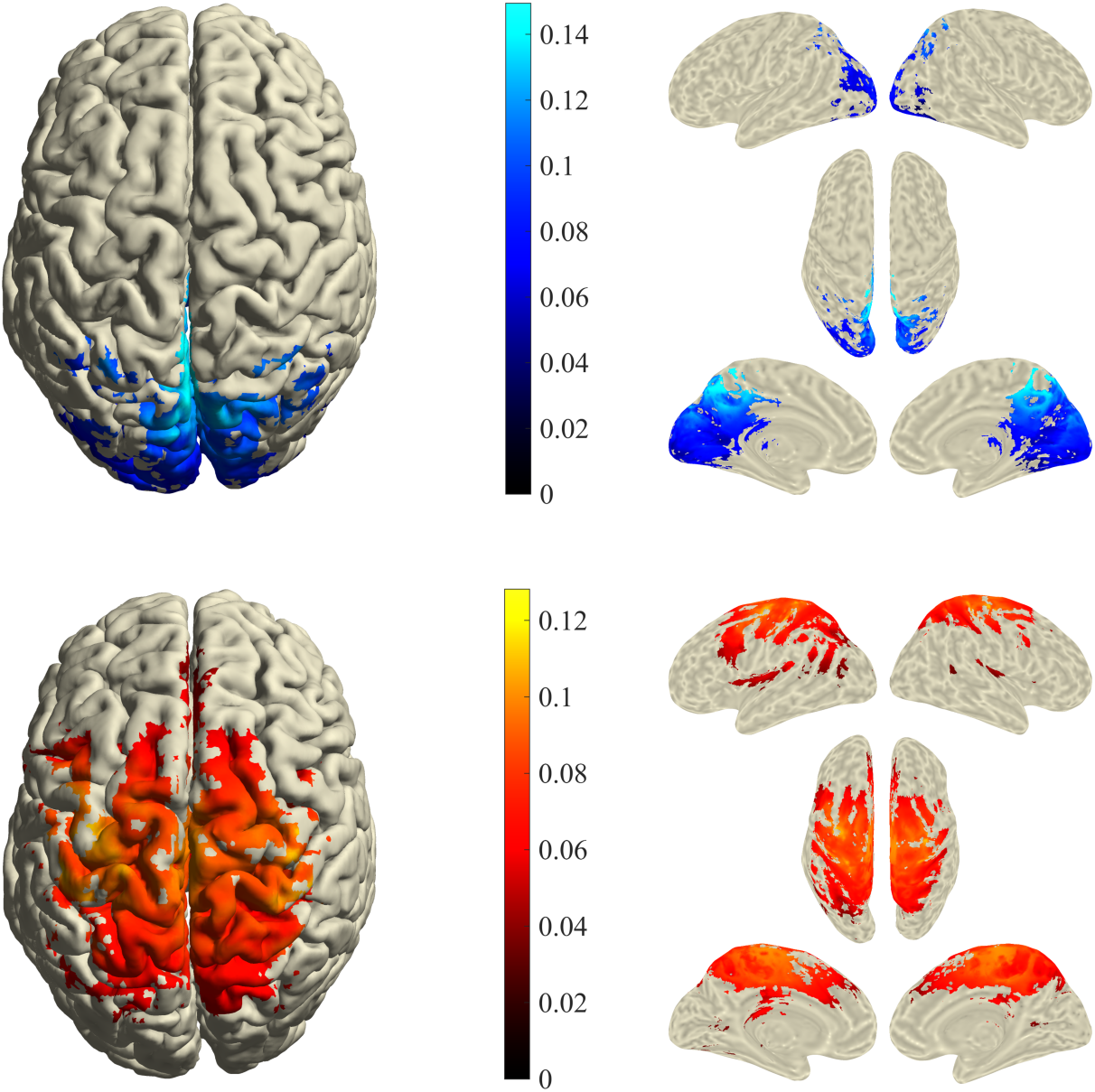
Power of the beamformer statistics masked for significance (values are normalized and, hence, units are arbitrary). The top row shows the main effect of age on pseudo-*t*-values (younger vs. older adults) in the alpha band. Significant voxels are mostly located in the cuneus and the precuneus. The bottom row displays the main effect of task-conditions (all task against rest/zero) in the beta band, and significant voxels predominantly located around the precentral gyrus, the superior frontal gyrus, and the juxtapositional lobule cortex. See Figure 4 for agreement with the selected ROIs. In Figure B.1 we provide the unmasked versions and in Figures B.2 and B.3 selected MRI slices including the corresponding MNI coordinates.

#### General statistical assessments

We adopted a 2×2×2 ANOVA design, where one factor was always *age* (younger vs. older adults) and the other two factors depended on the task/condition (see Table 2). Since we are dealing with repeated measures and mixed effects (fixed and random effects), we implemented linear mixed-effect (LME) models, which are not significantly influenced by outliers and non-normally distributed dependent variables (<u>Schielzeth et al., 2020</u>). In the *Results* section, we report the ANOVA marginal tests (of the fixed effects) and refer to the *Supplementary Material* for further details.

**Table 2.** 2×2 ANOVA tables for the two types of tasks (unimanual, panel A, and bimanual, panel B) with four conditions for each (RN, LN, RD and LD for unimanual task; SN, SD, ASRD and ASLD for bimanual task); see Table 1 for the definition of the abbreviations. The remaining factor *age* (younger vs. older) for the 2×2×2 ANOVA design is not shown.

| A |  | *hand side* |  |
| --- | --- | --- | --- |
|  |  | left | right |
| *hand difficulty level* | normal | LN | RN |
|  | difficult | LD | RD |

| B |  | *left-hand difficulty level* |  |
| --- | --- | --- | --- |
|  |  | normal | difficult |
| *right-hand difficulty level* | normal | SN | ASLD |
|  | difficult | ASRD | SD |

For the unimanual tasks, the remaining two fixed effect factors were the *hand difficulty level* (*HDL*: normal vs. difficult) and the *hand side* (*HS*: left hand vs. right hand). For the bimanual tasks, the fixed effects factors were the *left-hand difficulty level* (*LHDL*: normal vs. difficult) and the *right-hand difficulty level* (*RHDL*: normal vs. difficult). We also considered the two- and three-way interactions. The corresponding models can be formulated in Wilkinson notation as:

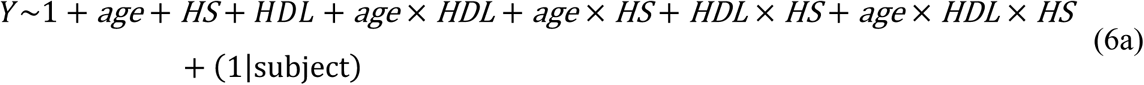

and

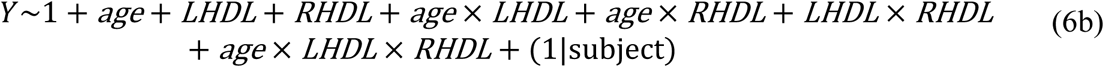

for the unimanual and bimanual cases, respectively, with *Y* being ℰ or *P̅*_ROI_ or *t̅*_ROI_ – see below – and participants were considered as a random effect. In Table 2 (A&B) we illustrate how each level relates to the task-conditions.

We adopted the same statistical design for the assessment of behavioral data and cortical activity. The behavioral data were evaluated via three LME models with different outcome variables: the total error: ℰ_Euclidean_, the left-hand error: 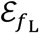, and the right-hand error: 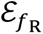. We averaged error values across the static and dynamic phases of performance. While ℰ_Euclidean_ served in both the unimanual and the bimanual tasks 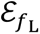 and 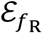 were only evaluated in the bimanual tasks to assess the left/right behavioral interference. For the alpha/beta power in the selected ROIs, we used the mean power *P̅*_ROI_ and the pseudo-*t*-value *t̅*_ROI_ that were computed from the virtual signals reconstructed by the respective spatial filters. For the alpha frequency band, we employed two LME models, one for (pre-)cuneus_L_ and one for (pre-)cuneus_R_. For the beta frequency band, we used four models, one for each (pre-)motor region: M1_L_, M1_R_, PMC_L_, and PMC_R_.

Finally, to identify associations between motor behavior and the spectral differences in the alpha and beta frequency band, we employed a similar LME design. We considered motor performance (ℰ_Euclidean_) as an output and the spectral outcome variables *X* (either *P̅*_ROI_ or *t̅*_ROI_) as predictors (fixed effect) along with *age* and task dependent factors that we already discussed for unimanual and bimanual tasks. Here, we only considered two-way interactions. The corresponding LME models read as follows:

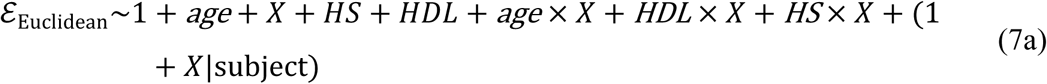

and

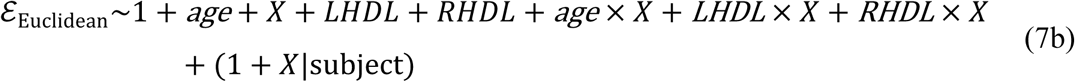

We would like to note that in contrast to the models (6a/b) both participants and the spectral values *X* were considered as random effects to capture subject-specific variations in the association between brain activity and performance error.

For every ROI, the LME models were fitted by restricted maximum likelihood estimation, and the normality of the residuals was verified via Shapiro-Wilks tests after visually inspecting the histograms and the Q-Q plots. For the *F*-tests, we employed Satterthwaite’s method to correct for sample bias (<u>Satterthwaite, 1946</u>). All *p*-values were corrected for multiple testing (over the number of ROIs per frequency) using the false discovery rate (FDR) correction (<u>Benjamini and Hochberg, 1995</u>; <u>Carvajal-Rodríguez et al., 2009</u>). Significant main or interaction effects were followed by *t*-tests for post hoc comparisons (independent in the case of age effects and paired in all other cases), for which we applied a Kenward-Roger approximation to estimate the denominator degrees-of-freedom. We applied a Bonferroni correction for multiple comparisons of the post hoc tests (<u>Armstrong, 2014</u>; <u>Chen et al., 2017</u>; <u>Noble, 2009</u>).

We performed the statistical assessments using RStudio, version 2023.6.1+524, with the lme4, lmerTest, emmeans, SGoF, and ggplot2 packages (<u>Bates et al., 2015</u>).

## Results

One participant from the younger and two from the older group had to be discarded from further analyses. One older adult pressed two sensors during the unimanual tasks and in two participants too many trials failed to pass our EEG preprocessing and data cleaning (one young adult had no data left for the RD task, one older adult had only two trials left for the ASRD task). Data from 18 younger and 17 older adults entered our analyses. Due to technical issues, three older adults could only perform the first two sets of trials containing 14 blocks in total (20 trials for each bimanual condition and 10 trials for each unimanual condition; one of them had 20 trials for the LD task). For the other participants, 30 trials for each bimanual condition and 20 trials for each unimanual condition were available.

In the subsequent subsection, we summarize our findings of the source localization when pooling all task-conditions before detailing task-specific effects at both behavioral and cortical levels including the associations between cortical activity and behavioral performance.

### Source localization

#### Alpha activity

The beamformer analysis in the alpha frequency band revealed a significant main effect of *age* for the power contrast between the dynamic and the static phases (pseudo-*t*-value) in parieto-occipital areas (Figure 5).

We also found a significant main effect of task in the same brain regions. For the sake of legibility, however, we here focus on the age-related regions and refer to Figure S.2 in the *Supplementary Material* for the task-related beamformer results. To illustrate the activity in these areas, we provide in Figure 6 the wavelet spectra at source level that were estimated in line with our pre-analysis (cf. Figure 3).

**Figure 6.**
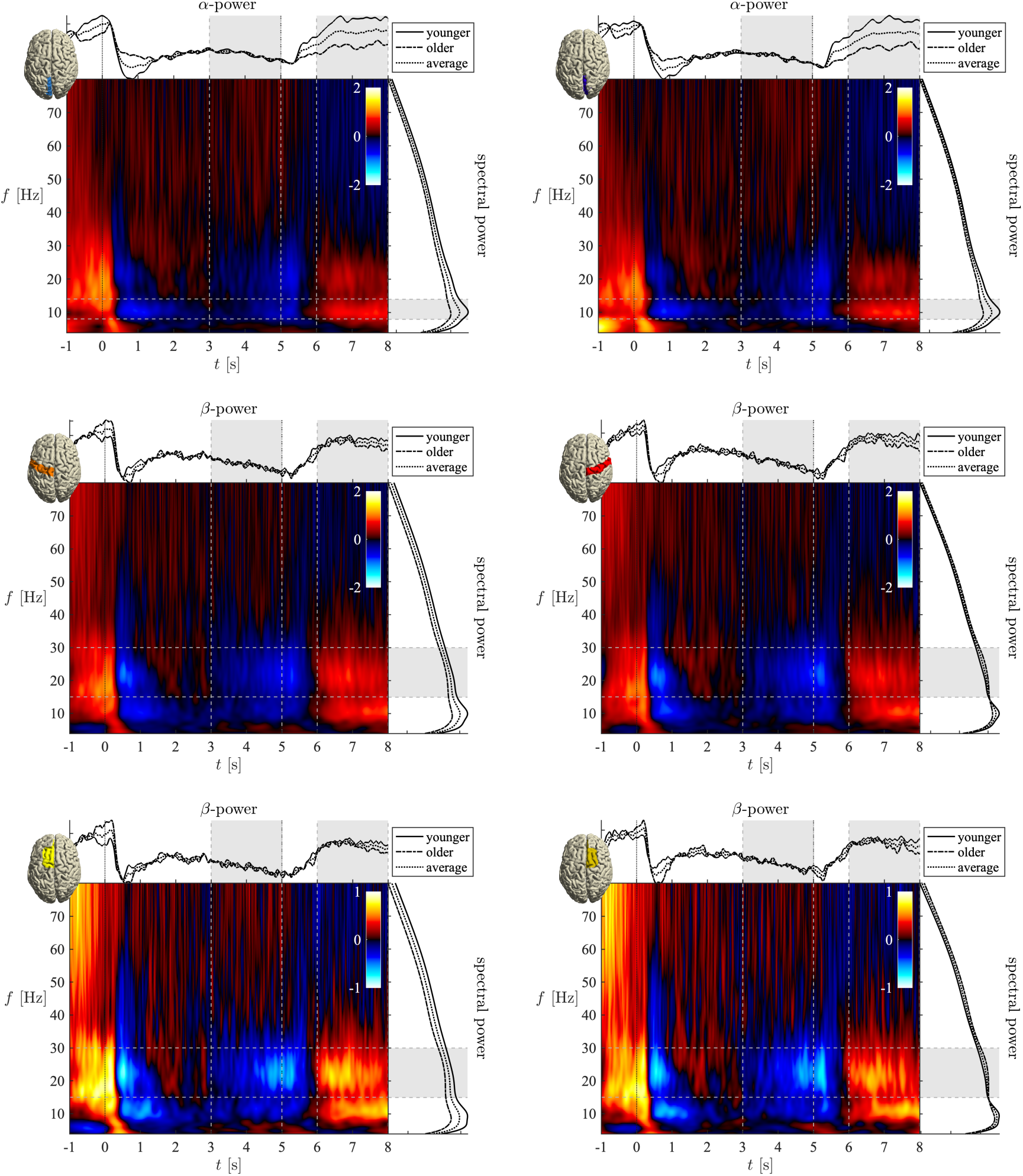
Top row: Spectra of bilateral visual areas. Left panel = (pre)cuneus_L_; right panel = (pre)cuneus_R_. We computed the average over trials per task-condition, averaged all task-conditions per participant, and finally averaged over all participants within each age group. The time-frequency spectra are the averages over all participants of both age groups (normalized per frequency to the [1-8]s interval). Second and third rows: Wavelet spectra of bilateral M1 and PMC regions (2^nd^ row, left/right panels: M1_L/R_; 3^rd^ row left/right panels: PMC_L/R_). Power spectra (2D lines, right to the wavelet spectra) and time-dependent power (2D lines, on the top of the wavelet spectra) are group-specific. The latter were normalized to the [3,5]s = ERD interval to ease visual inspection and highlight age-related differences. Gray bars indicate the epochs and the frequency bands of interest; cf. Figure 3.

As expected, the most visible power differences at source level could be found in the alpha frequency band, for which not only the total power differed between age groups, but especially the ERS power during the static phase was clearly lower in the older participants than in the younger ones in this visualization, at least relative to the respective ERD power in the dynamic phase (Figure 6 top panels).

#### Beta activity

In the beta frequency band, the pseudo-*t*-values reached significance over the motor network for the main effect of task. In contrast to the alpha activity, however, we could not establish a main effect of *age* in the beta band. The lower panels in Figure 5 depict the *t*-statistic related to the main effect of task performance on beta power modulations across participants. These results clearly overlap with the ROIs depicted in Figure 4, namely bilateral precentral gyrus (left/right M1) and bilateral superior frontal gyrus (left/right PMC, including SMA), which confirms the appropriateness of our choice to mask the spatial filters via the respective regions of Harvard/Oxford cortical atlas when projecting the EEG to the sources.

For the corresponding time-frequency plots, we refer to Figure 6. Beta-ERD/ERS cycles are clearly visible in all regions, corresponding to the dynamic and static phases of the motor tasks, respectively. As for the alpha activity, we found significant age effects in the beta activity for the mean power over time (see the 2D lines on the right side of the images in Figure 6), but after normalizing to the mean power in the 3-5s interval, this effect was not observed in beta power, as it was in alpha power.

### Unimanual conditions

The ANOVAs for the performance error, the mean power, and the pseudo-*t*-value for each ROI revealed multiple significant main and interaction effects that we summarized in Table 3. The subsequent paragraphs highlight the most important outcomes, first for the behavioral data, followed by the EEG data. All the reported post hoc results reached significance, see the *Supplementary Material* for more details on mean and *p*-values.

**Table 3.** ANOVA results based on the linear mixed-effects model *Y* ∼ 1 + *age* × *HDL* + *age* × *HS* + *HDL* × *HS* + *age* × *HDL* × *HS* + (1 | subject) for *Y* being the performance error (ℰ), the mean power (*P̅*) or the pseudo-*t*-value (*t̅*) in left/right (pre-)cuneus (alpha) and in M1_L_, M1_R_, PMC_L_, and PMC_R_ (beta) during the unimanual tasks.*

| *Y* | *age* |  | *HDL* |  | *HS* |  | *age* × *HDL* |  | *age* × *HS* |  | *HDL* × *HS* |  | *age* × *HDL* × *HS* |  |
| --- | --- | --- | --- | --- | --- | --- | --- | --- | --- | --- | --- | --- | --- | --- |
|  | *F*<sub>33,1</sub> | *p* | *F*<sub>99,1</sub> | *p* | *F*<sub>99,1</sub> | *p* | *F*<sub>99,1</sub> | *p* | *F*<sub>99,1</sub> | *p* | *F*<sub>99,1</sub> | *p* | *F*<sub>99,1</sub> | *p* |
| $\mathcal{E}_{\text{Euc}}$ | 13.6 | <10<sup>-3</sup> | 838.6 | <10<sup>-3</sup> | 0.2 | .672 | 1.2 | .272 | 0.7 | .421 | 0.2 | .666 | 0.2 | .655 |
| $\bar{P}_{\text{preCL}}(\alpha)$ | 20.1 | <10<sup>-3</sup> | 6.3 | .014 | 0.5 | .501 | 0.1 | .735 | 4.4 | .039 | 6.7 | .022 | 0.1 | .937 |
| $\bar{P}_{\text{preCR}}(\alpha)$ | 14.6 | <10<sup>-3</sup> | 7.1 | .014 | 2.8 | .199 | 0.5 | .735 | 5.8 | .036 | 3.3 | .071 | 0.0 | .937 |
| $\bar{t}_{\text{preCL}}(\alpha)$ | 16.6 | <10<sup>-3</sup> | 12.2 | .001 | 1.1 | .303 | 0.0 | .901 | 0.0 | .971 | 2.7 | .204 | 1.2 | .562 |
| $\bar{t}_{\text{preCR}}(\alpha)$ | 15.0 | <10<sup>-3</sup> | 8.0 | .006 | 1.8 | .303 | 1.1 | .606 | 1.0 | .631 | 1.1 | .305 | 0.0 | .860 |
| $\bar{P}_{\text{M1L}}(\beta)$ | 5.1 | .062 | 2.7 | .424 | 8.5 | .017 | 1.3 | .348 | 6.4 | .013 | 9.7 | .005 | 0.3 | .991 |
| $\bar{P}_{\text{M1R}}(\beta)$ | 0.5 | .479 | 0.0 | .956 | 3.6 | .079 | 0.4 | .555 | 6.8 | .013 | 7.0 | .009 | 0.0 | .991 |
| $\bar{t}_{\text{M1L}}(\beta)$ | 1.2 | .450 | 11.1 | .005 | 0.7 | .530 | 0.3 | .602 | 6.3 | .049 | 9.2 | .009 | 0.4 | .766 |
| $\bar{t}_{\text{M1R}}(\beta)$ | 1.0 | .450 | 9.7 | .005 | 19.0 | <10<sup>-3</sup> | 1.2 | .546 | 0.8 | .379 | 2.0 | .163 | 0.3 | .766 |
| $\bar{P}_{\text{PMCL}}(\beta)$ | 6.8 | .054 | 0.1 | .939 | 7.1 | .018 | 5.3 | .095 | 7.0 | .013 | 11.3 | .004 | 0.4 | .991 |
| $\bar{P}_{\text{PMCR}}(\beta)$ | 0.7 | .479 | 0.5 | .939 | 1.0 | .310 | 2.2 | .290 | 8.1 | .013 | 8.4 | .006 | 0.0 | .991 |
| $\bar{t}_{\text{PMCL}}(\beta)$ | 0.6 | .450 | 7.7 | .009 | 0.4 | .530 | 0.3 | .602 | 5.2 | .049 | 8.4 | .009 | 0.5 | .766 |
| $\bar{t}_{\text{PMCR}}(\beta)$ | 0.6 | .450 | 4.3 | .041 | 10.5 | .003 | 1.3 | .546 | 1.9 | .230 | 2.5 | .152 | 0.0 | .871 |

\* *HDL* = hand difficulty level, *HS* = hand side; all *p*-values are FDR adjusted; bold-face indicates statistical significance.

#### Behavioral data

We found significant main effects of *age* and *hand difficulty level*, while neither a significant difference between left- and right-hand performance (*hand side*) nor any significant interactions were found (top row in Table 3). Overall, older adults showed larger performance errors than younger adults, as expected. As shown in Figure 7, in both age groups the errors increased with increasing task difficulty.

**Figure 7.**
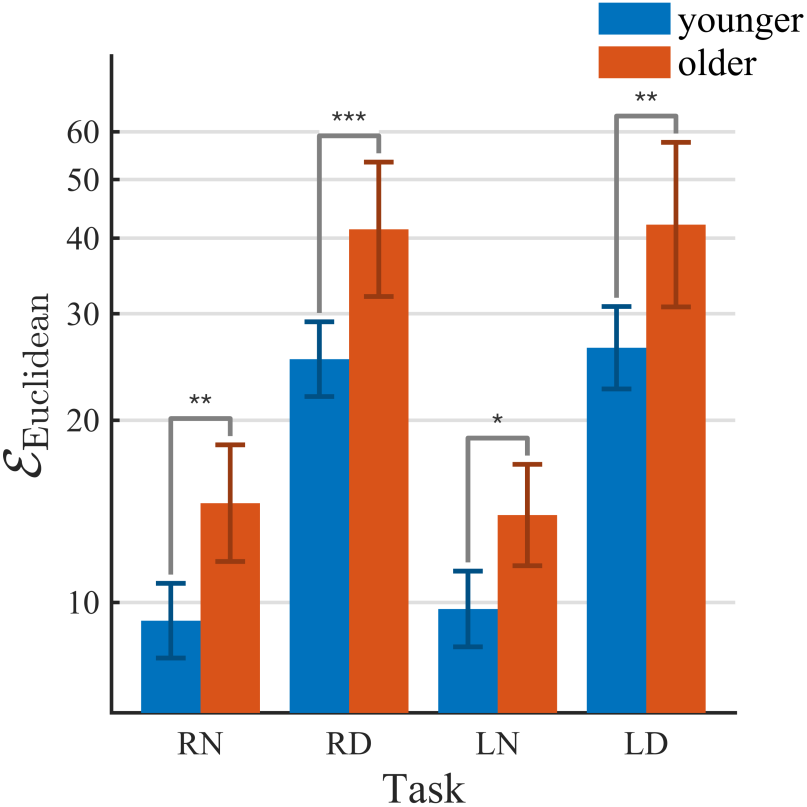
Average values of the performance errors (ℰ_Euclidean_) during the unimanual tasks. In the more challenging tasks (RD and LD), the error was significantly higher than in the normal conditions (RN and LN) and more specifically for older adults, albeit without significant interaction effect; error bars reflect the 95% confidence interval (CI) and significant between-group effects (corrected for multiple comparisons) are indicated using *\** ∼ *p*<.05, ** ∼ *p*<.01, *** ∼ *p*<10^-3^, and **** ∼ *p* < 10^-4^.

#### Alpha activity

Like the behavioral data, the mean power and pseudo-*t*-values (normalized difference between ERS and ERD power) in left/right (pre-)cuneus revealed significant main effects of *age* and *hand difficulty level,* but not of *hand side*. Only for the mean power, some selected interactions reached significance (see Table 3, 2^nd^ to 5^th^ row). Older adults showed a lower alpha mean power and power modulation (i.e., normalized contrast between ERS and ERD = pseudo-*t*-value) than the younger ones and both measures were higher in normal than in difficult conditions. Especially, the mean power in (pre-)cuneus_L_ was higher in the LN than in the LD condition; see Figure 8 and Table S.2 for further details.

**Figure 8.**
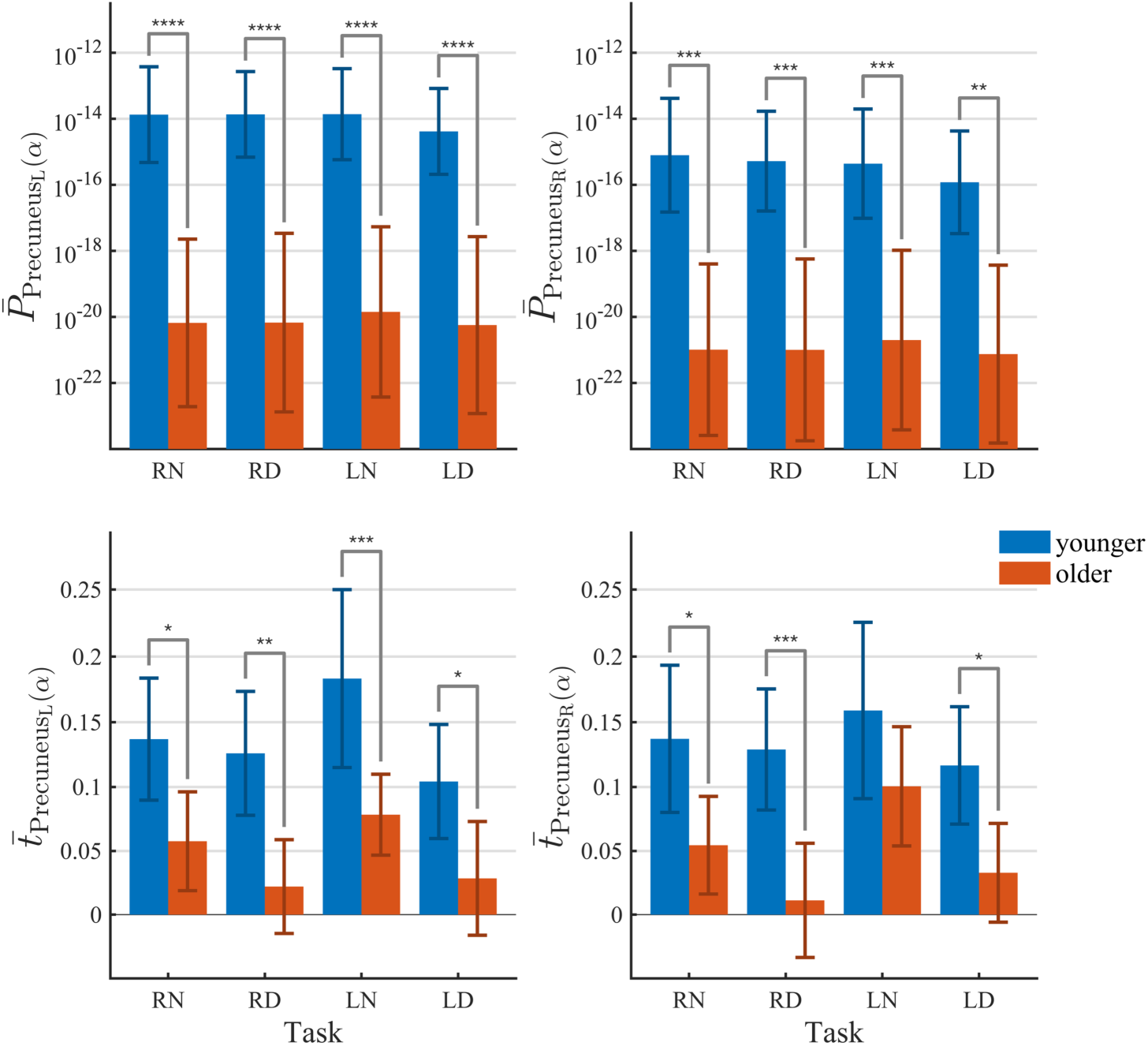
Average values of mean power *P̅*(*α*) and pseudo-*t*-values *t̅*(*α*) in precuneus_L/R_ during the unimanual tasks. In the more challenging tasks (RD and LD), the power modulation was significantly lower than in the normal conditions (RN and LN) and more specifically for older adults. Error bars indicate the 95% CI and significant between-group effects (corrected for multiple comparisons) are indicated using * ∼ *p*<.05, ** ∼ *p*<.01, *** ∼ *p*<10^-3^, and **** ∼ *p* < 10^-4^.

As for the significant *age* × *hand side* interaction effect in bilateral (pre-)cuneus, post-hoc tests always revealed a lower mean power in older compared to younger adults, but the age difference was more pronounced during right-hand compared to left-hand performance and only younger adults showed higher mean power during right-hand compared to left hand performance. We also found a significant *hand difficulty level* × *hand side* interaction for the left (pre-)cuneus, where the mean power in the normal condition was higher than in the difficult condition during left-hand performance only (Table S.3).

#### Beta activity

As shown in Table 3 (rows 6-13), there was a significant main effect of *hand difficulty level* across motor areas for the power modulation but not for the mean power. In all participants, the normal conditions were accompanied by larger power modulations than the ones during difficult conditions. The main effect of *hand side* for the mean power turned out significant only in PMC_L_ and M1_L_, and for the power modulation in M1_R_ as well as PMC_R_ (Figure 9). During left-hand performance, both mean power and power modulation were higher than during right-hand performance (Table S.4). We did not observe a significant main effect of *age*, however, there were significant interaction effects: (i) *age* × *hand side,* and (ii) *hand difficulty level* × *hand side* for the mean power across all regions, and for the power modulation in M1_L_ and PMC_L_. The mean power in older adults was always lower than in younger adults in M1_L_ and PMC_L_, and in M1_R_ and PMC_R_ the mean power was higher during right- than during left-hand performances only in the younger adults. In M1_R_ and PMC_R_, the older adults showed lower power modulations during right-hand compared to left-hand performance. With regards to the *hand difficulty level* × *hand side* interaction, in the normal condition, both mean power and power modulation during right-hand performances were lower than during left-hand performances in M1_L_ and PMC_L_, while in the difficult condition, the mean power was higher during right-hand compared to left-hand performances in M1_R_ and PMC_R_ (Table S.5).

**Figure 9.**
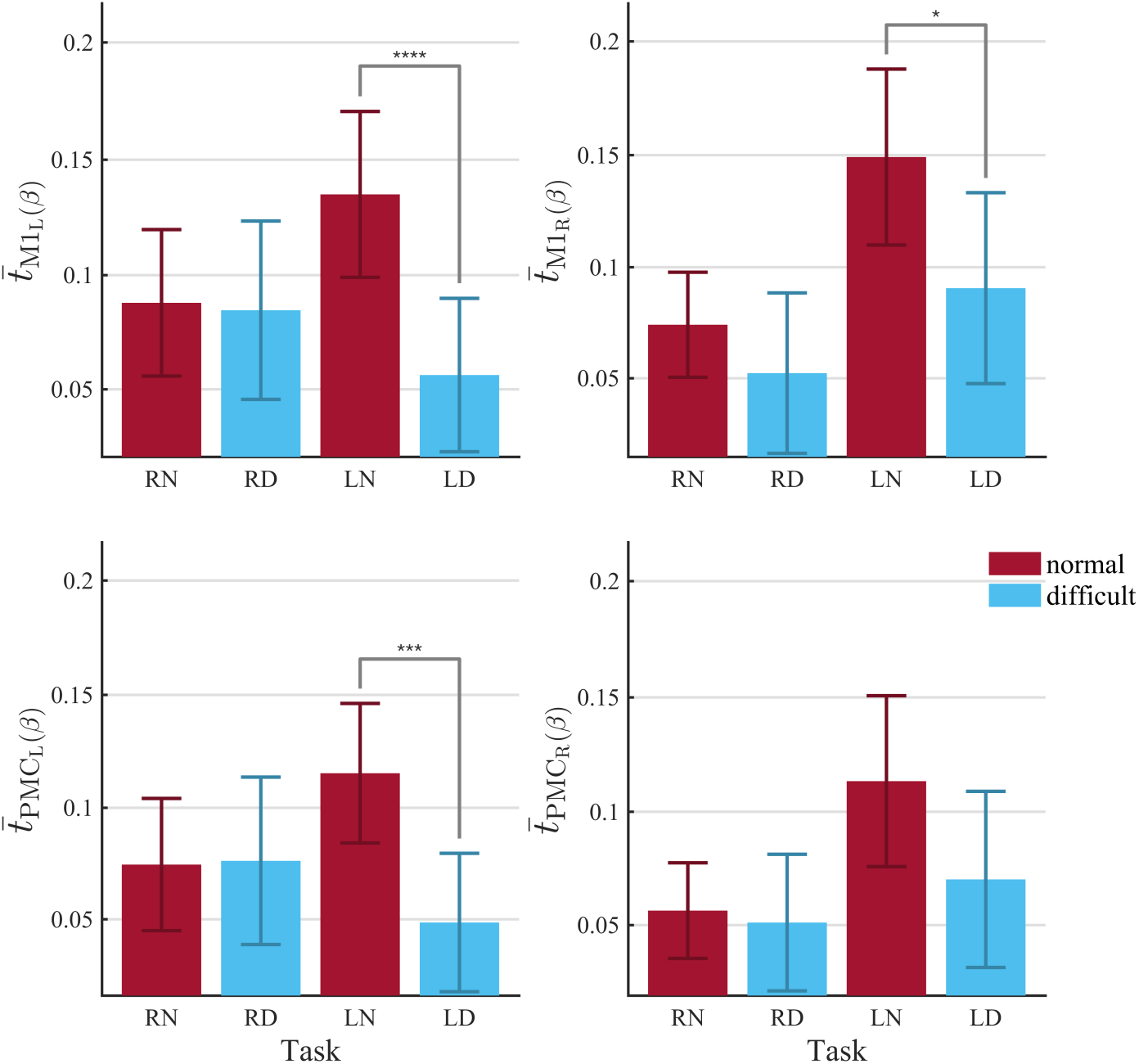
Average pseudo-*t*-values *t̅*(*β*) in M1_L/R_ and PMC_L/R_ during the unimanual tasks. In the more challenging tasks (RD and LD), the power modulation was significantly lower than in the normal conditions (RN and LN). Error bars show the 95% CI, significant within-group effects (corrected for multiple comparisons) are indicated using * ∼ *p*<.05, ** ∼ *p*<.01, *** ∼ *p*<10^-3^, and **** ∼ *p* < 10^-4^.

#### Association between error of performance and alpha/beta activity

As outlined in the *Statistics* section, we also incorporated linear mixed-effects models to estimate the associations between the error of performance and both mean power and power modulations in the alpha and beta frequency bands in (pre-)cuneus_L/R_, M1_L/R_, and PMC_L/R_. While the mean power did not show any significant association with ℰ_Euclidean_, we found significant effects for the power modulation. The results are presented in Table 4; see especially column *X* in that table. In the unimanual tasks, this association was limited to bilateral M1 and (pre-)cuneus_R_; see also **Error! Reference source not found.** (top panel) below.

**Table 4.**
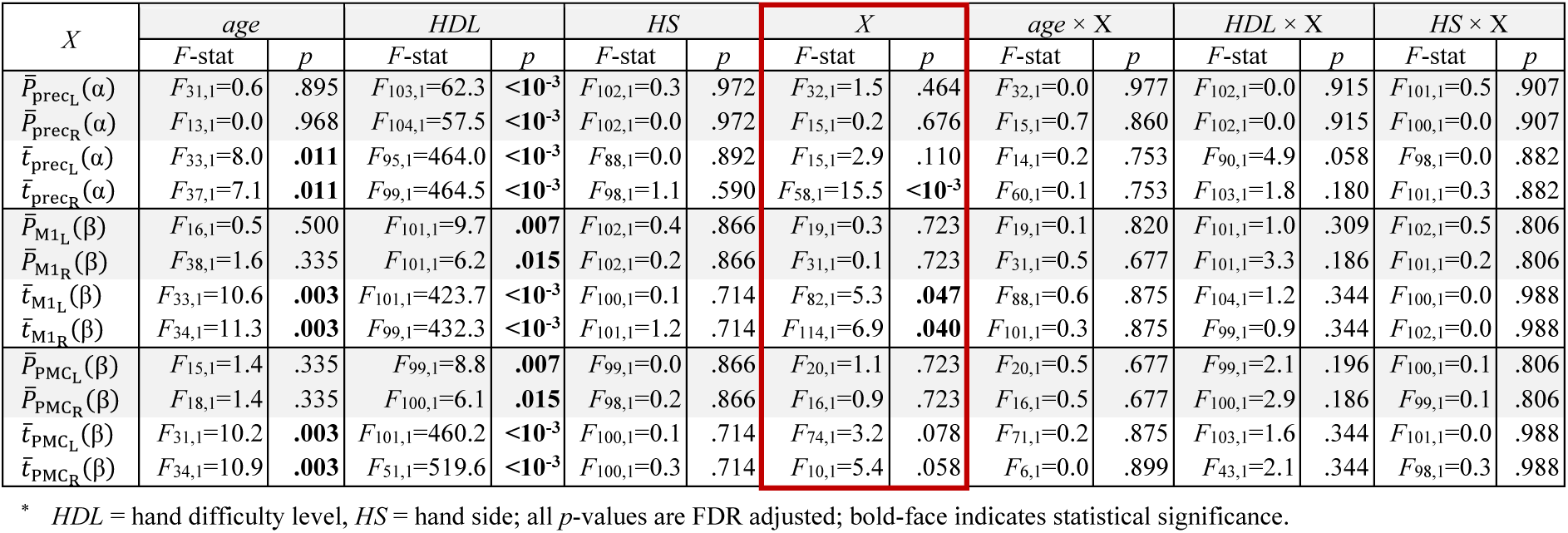
ANOVA results of the LME ℰ_Euc._ ∼ 1 + *age* × *X* + *HDL* × *X* + *HS* × *X* + (1 + *X* | subject) for *X* being the mean power (*P*\*), or the pseudo-*t*-value (*t̅*) in all ROIs during the unimanual tasks.*

### Bimanual conditions

Again, we first provide an overview of the ANOVAs outcomes in Table 5 and subsequently report only the post hoc effects that reached significance and refer to the *Supplementary Material* for further details.

**Table 5.** ANOVA results from the LME *Y* ∼ 1 + *age* × *LHDL* + *age* × *RHDL* + *LHDL* × *RHDL* + *age* × *LHDL* × *RHDL* + (1 | subject) for *Y* being the performance errors (ℰ), the mean power (*P*\*), or the pseudo-*t*-value (*t̅*) of (pre-)cuneus_L_, (pre-)cuneus_R_ (alpha activity), M1_L_, M1_R_, PMC_L_, and PMC_R_ (beta activity) during the bimanual tasks.*

| $Y$ | $age$ | | $LHDL$ | | $RHDL$ | | $age \times LHDL$ | | $age \times RHDL$ | | $LHDL \times RHDL$ | | $age \times LHDL \times RHDL$ | |
| --- | --- | --- | --- | --- | --- | --- | --- | --- | --- | --- | --- | --- | --- | --- |
| | $F_{33,1}$ | $p$ | $F_{99,1}$ | $p$ | $F_{99,1}$ | $p$ | $F_{99,1}$ | $p$ | $F_{99,1}$ | $p$ | $F_{99,1}$ | $p$ | $F_{99,1}$ | $p$ |
| $\mathcal{E}_{Eucl}$ | 17.2 | **<10<sup>-3</sup>** | 230.8 | **<10<sup>-3</sup>** | 269.0 | **<10<sup>-3</sup>** | 0.0 | .922 | 0.0 | .991 | 522.4 | **<10<sup>-3</sup>** | 0.1 | .747 |
| $\mathcal{E}_{f_L}$ | 11.8 | **.002** | 152.1 | **<10<sup>-3</sup>** | 26.2 | **<10<sup>-3</sup>** | 0.7 | .591 | 1.8 | .552 | 145.4 | **<10<sup>-3</sup>** | 3.8 | .126 |
| $\mathcal{E}_{f_R}$ | 13.1 | **.001** | 34.8 | **<10<sup>-3</sup>** | 245.0 | **<10<sup>-3</sup>** | 5.0 | .083 | 0.0 | .991 | 177.6 | **<10<sup>-3</sup>** | 3.1 | .126 |
| $\bar{P}_{preCL}(\alpha)$ | 18.4 | **<10<sup>-3</sup>** | 1.0 | .504 | 3.3 | .073 | 0.0 | .838 | 0.4 | .537 | 8.2 | **.010** | 2.2 | .137 |
| $\bar{P}_{preCR}(\alpha)$ | 12.7 | **.001** | 0.4 | .504 | 3.8 | .073 | 0.2 | .838 | 0.5 | .537 | 4.8 | **.031** | 4.1 | .092 |
| $\bar{t}_{preCL}(\alpha)$ | 19.3 | **<10<sup>-3</sup>** | 0.0 | .971 | 2.7 | .103 | 2.2 | .291 | 1.6 | .416 | 7.3 | **.013** | 0.2 | .647 |
| $\bar{t}_{preCR}(\alpha)$ | 22.7 | **<10<sup>-3</sup>** | 0.1 | .971 | 4.8 | .061 | 1.0 | .318 | 0.7 | .420 | 6.4 | **.013** | 0.3 | .647 |
| $\bar{P}_{M1L}(\beta)$ | 5.4 | .054 | 8.5 | **.004** | 1.5 | .231 | 4.3 | .080 | 3.4 | .193 | 1.3 | .760 | 0.4 | .949 |
| $\bar{P}_{M1R}(\beta)$ | 0.5 | .482 | 10.5 | **.004** | 1.9 | .231 | 3.0 | .112 | 1.4 | .247 | 0.3 | .760 | 0.1 | .949 |
| $\bar{t}_{M1L}(\beta)$ | 0.5 | .602 | 2.0 | .265 | 10.9 | **.004** | 0.4 | .519 | 0.0 | .970 | 9.5 | **.011** | 0.0 | .895 |
| $\bar{t}_{M1R}(\beta)$ | 1.4 | .534 | 2.5 | .265 | 6.4 | **.013** | 0.6 | .519 | 0.2 | .943 | 6.2 | **.019** | 0.0 | .895 |
| $\bar{P}_{PMC_L}(\beta)$ | 7.2 | **.044** | 8.5 | **.004** | 3.4 | .231 | 2.5 | .114 | 2.8 | .193 | 0.8 | .760 | 0.0 | .995 |
| $\bar{P}_{PMC_R}(\beta)$ | 0.6 | .482 | 9.2 | **.004** | 2.4 | .231 | 4.4 | .080 | 2.0 | .216 | 0.0 | .905 | 1.1 | .949 |
| $\bar{t}_{PMC_L}(\beta)$ | 0.3 | .602 | 1.7 | .265 | 10.0 | **.004** | 1.0 | .519 | 0.1 | .943 | 8.0 | **.012** | 0.0 | .895 |
| $\bar{t}_{PMC_R}(\beta)$ | 1.3 | .534 | 0.7 | .395 | 8.4 | **.006** | 1.0 | .519 | 1.3 | .943 | 2.4 | .124 | 0.2 | .895 |

\* $LHDL$ = left-hand difficulty level, $RHDL$ = right-hand difficulty level; all $p$ -values are adjusted via FDR correction; bold-face indicates statistical significance.

#### Behavioral data

As expected, the ANOVAs of the LME model’s marginals revealed significant main effects of *age*, *left- and right-hand difficulty level,* and a significant interaction effect between *left- and right-hand difficulty level* for all performance measures. Yet, neither the *age* × *left/right-hand difficulty level,* nor the three-way interaction reached significance (Table 5, top three rows). Recall that ℰ_Euclidean_indicates the error of performance on the screen, whereas 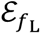 and 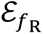 indicate the error of left- and right-hand force productions, respectively. As shown in Figure 10, older adults performed all bimanual tasks less accurately than younger ones. The performance errors were generally larger with a higher degree of difficulty (Table S.6). When the *left-hand difficulty level* was normal, the error of performance was higher when the *right-hand difficulty level* was high (ASRD) than when it was normal (SN) and vice versa; cf. Table S.7. Importantly, the results for 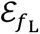 and 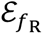 confirm that the left-hand difficulty level induced by amplifying the high-frequency tremor does affect the performance of the right hand and vice versa, which is in line with our corresponding hypothesis.

**Figure 10.**
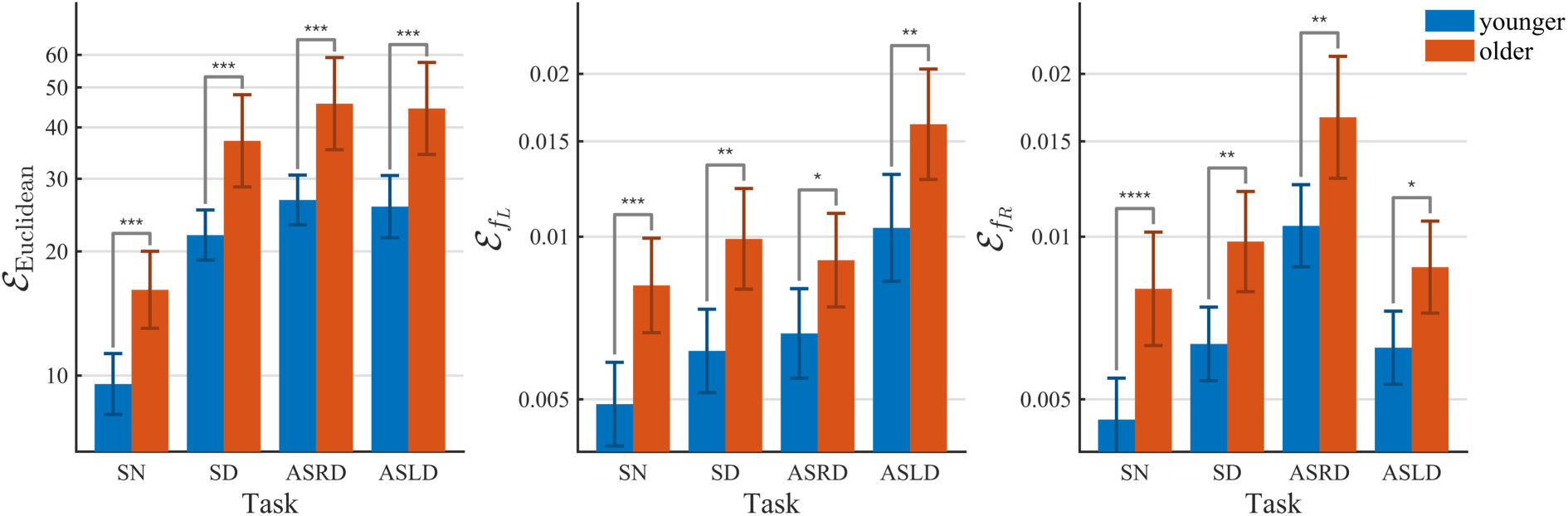
Average values of the performance errors during the bimanual task. In the more challenging tasks (ASRD and ASLD), the errors were significantly higher than in the symmetric normal one (SN); error bars indicate the 95% CI and significant between-group effects (corrected for multiple comparisons) are indicated using * ∼ *p*<.05, ** ∼ *p*<.01, *** ∼ *p*<10^-3^, and **** ∼ *p* < 10^-4^.

#### Alpha activity

For the mean power and the power modulation in left/right (pre-)cuneus, we found a significant main effect of *age* and a significant *left-* × *right-hand difficulty level* interaction (Table 5, rows 4-7) with older adults showing lower mean power and pseudo-*t*-values; see Figure 11 and Table S.8.

**Figure 11.**
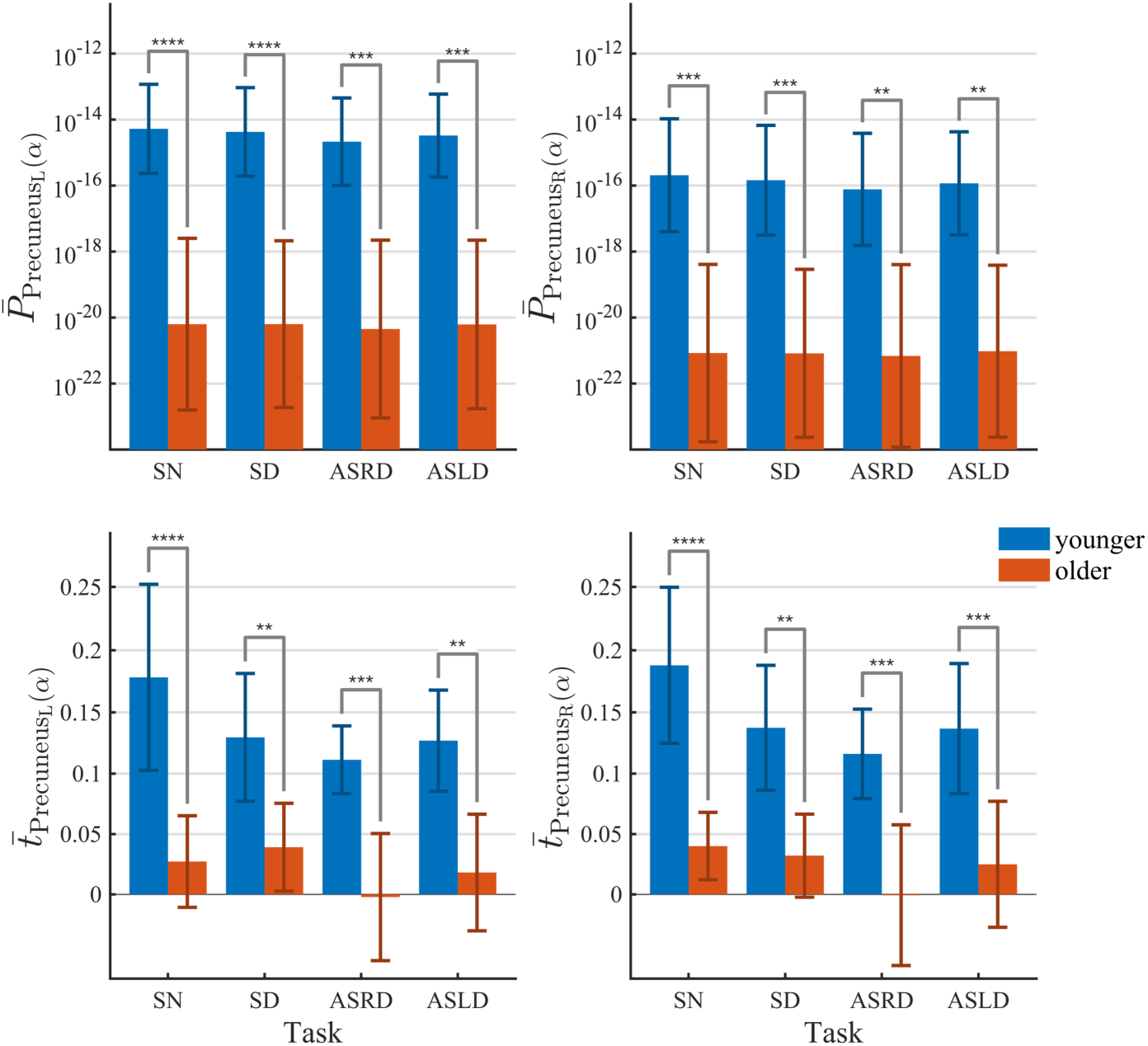
Average values of the mean power *P̅*(*α*) and pseudo-*t*-value *t̅*(*α*) in bilateral (pre-)cuneus during the bimanual task in the younger and older groups. In ASRD condition, the mean power and power modulation were lower than in other conditions; ASRD task displayed the lowest pseudo-*t*-value. Error bars indicate the 95% CI, significant effects (corrected for multiple comparisons) are indicated via * ∼ *p*<.05, ** ∼ *p*<.01, *** ∼ *p*<10^-3^, and **** ∼ *p* < 10^-4^.

In the normal left-hand conditions (SN & ASRD), the mean power and modulation were higher when the *right-hand difficulty level* was normal (i.e., SN). That is, both measures were lower in the ASRD task compared to the SN task (Table S.9).

#### Beta activity

As shown in Table 5 (rows 8-15), there was a significant main effect of *age* for the mean beta power in PMC_L_ as well as a significant main effect of *left-* and *right-hand difficulty level* for both mean power and power modulation, respectively (Figure 12). We also found a significant *left-* × *right-hand difficulty level* interaction for the power modulation in M1_L_, M1_R_ and PMC_L_. All other main and interaction effects did not reach significance.

**Figure 12.**
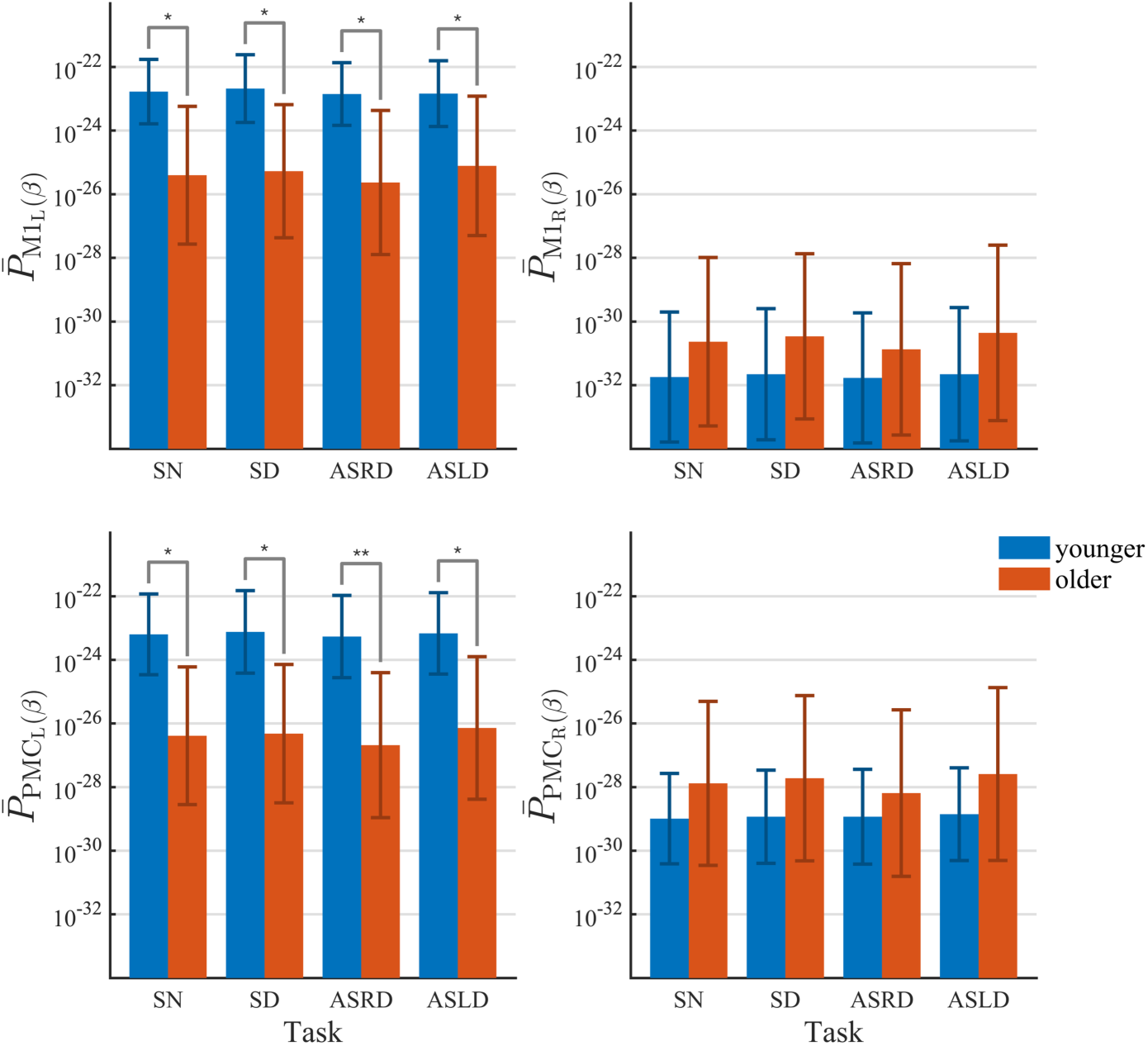
Average values of the mean power *P̅*(*β*) in M1_L/R_ and PMC_L/R_ during the bimanual task in the younger and older groups. Error bars show the 95% CI and significant effects (corrected for multiple comparisons) are indicated using * ∼ *p*<.05, ** ∼ *p*<.01, *** ∼ *p*<10^-3^, and **** ∼ *p* < 10^-4^

**Figure 13.**
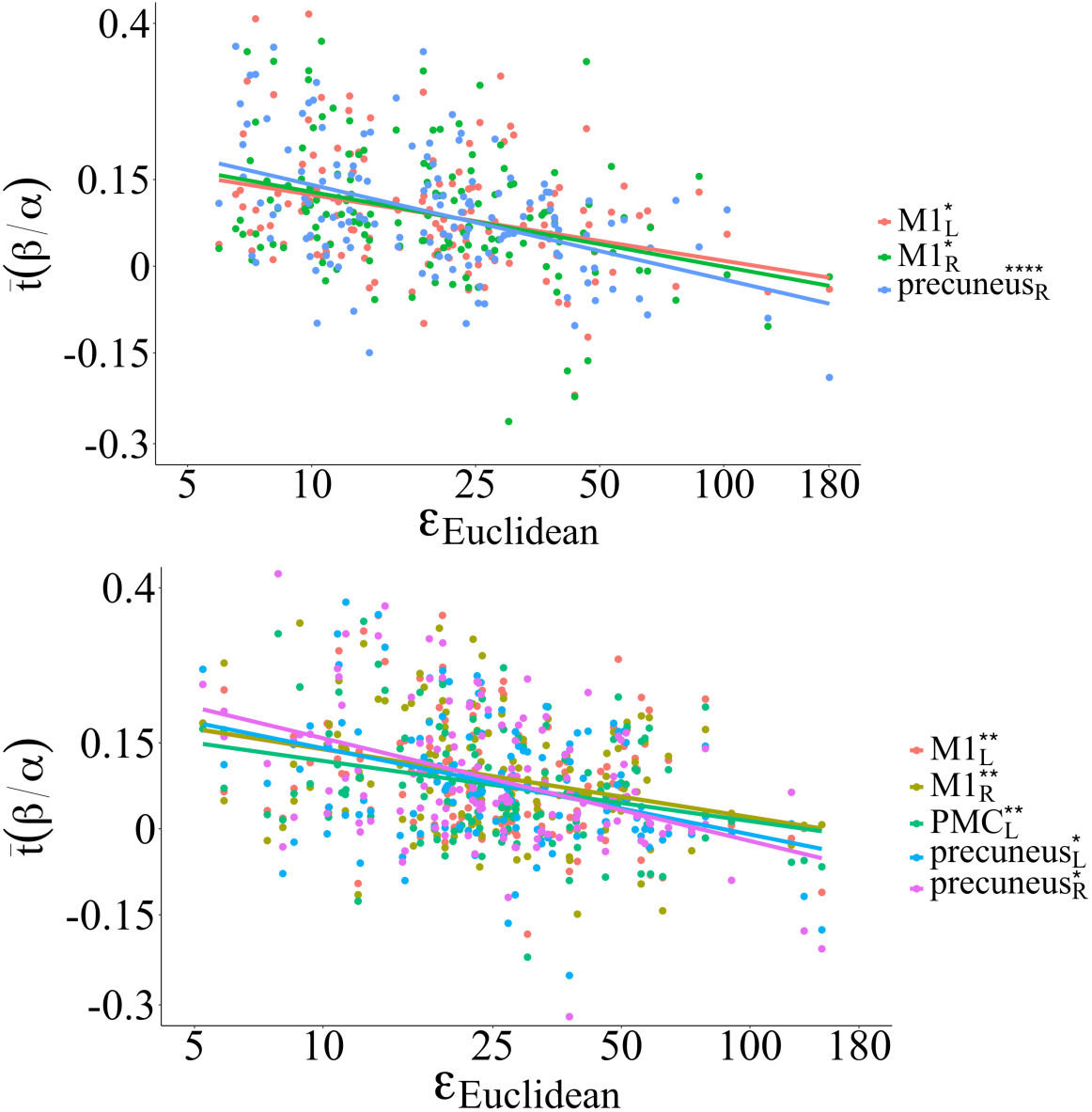
Associations between the performance error and the beta modulation (pseudo-*t*-value) during the unimanual (top panel) and bimanual task (bottom panel) across groups and conditions. The scatter plot indicates that large beta power modulation (higher pseudo-*t*-values) are associated with better quality of performance. Significant associations (corrected for multiple comparisons) are indicated in the legends using * ∼ *p*<.05, ** ∼ *p*<.01, *** ∼ *p*<10^-3^, and **** ∼ *p* < 10^-4^; cf. Tables 5 & 6.

In PMC_L_, the mean beta power in older adults was significantly lower than in younger ones. For all motor regions, both groups performed the left-hand normal conditions (SN & ASRD) with lower mean power than the left-hand difficult conditions (SD & ASLD). However, right-hand performance was accompanied by a larger power modulation in the normal than in the difficult conditions (see Table S.10). In M1_L_, M1_R_, and PMC_L_, the power modulation was lower when the condition of one hand differed from the other hand than when the condition of both hands was normal. E.g., when the left-hand difficulty level was normal, pseudo-*t*-values were lower in the difficult condition for the right (ASRD) hand compared to the normal condition (SN; cf. Table S.11).

Interestingly, Figure 12 also suggests a hemispheric difference in that, across conditions, the age-related reduction in mean power and power modulation in the left hemisphere is ‘compensated’ by an increase in the right hemisphere. Although the latter did not reach significance, we will return to this in de *Discussion* section.

#### Association between error of performance and alpha/beta activity

The results of the linear mixed-effects models for the associations between the error of performance and both the mean power and the power modulation in the alpha and beta frequency bands in the different ROIs are summarized in Table 6 (cf. column *X*).

**Table 6.** ANOVA results for the LME ℰ_Euc._ ∼ 1 + *age* × *X* + *LHDL* × *X* + *RHDL* × *X* + (1 + *X* | subject) for *X* being the mean power (*P*\*), or the pseudo-*t*-value (*t̅*) in all ROIs during the bimanual tasks.*

| $X$ | *age* | | *LHDL* | | *RHDL* | | $X$ | | *age* × $X$ | | *LHDL* × $X$ | | *RHDL* × $X$ | |
| --- | --- | --- | --- | --- | --- | --- | --- | --- | --- | --- | --- | --- | --- | --- |
|  | *F*-stat | *p* | *F*-stat | *p* | *F*-stat | *p* | *F*-stat | *p* | *F*-stat | *p* | *F*-stat | *p* | *F*-stat | *p* |
| $\bar{P}_{\text{precL}}(\alpha)$ | $F_{27,1}=3.2$ | .169 | $F_{101,1}=5.4$ | **.031** | $F_{100,1}=1.6$ | .213 | $F_{29,1}=2.4$ | .267 | $F_{29,1}=1.1$ | .608 | $F_{101,1}=0.4$ | .536 | $F_{100,1}=0.3$ | .649 |
| $\bar{P}_{\text{precR}}(\alpha)$ | $F_{14,1}=1.4$ | .250 | $F_{100,1}=4.8$ | **.031** | $F_{100,1}=1.6$ | .213 | $F_{15,1}=0.7$ | .425 | $F_{15,1}=0.1$ | .710 | $F_{100,1}=0.4$ | .536 | $F_{100,1}=0.2$ | .649 |
| $\bar{t}_{\text{precL}}(\alpha)$ | $F_{36,1}=4.9$ | .051 | $F_{103,1}=25.7$ | **<10<sup>-3</sup>** | $F_{102,1}=35.4$ | **<10<sup>-3</sup>** | $F_{69,1}=5.2$ | **.028** | $F_{69,1}=0.2$ | .681 | $F_{107,1}=0.0$ | .985 | $F_{109,1}=1.3$ | .355 |
| $\bar{t}_{\text{precR}}(\alpha)$ | $F_{38,1}=4.1$ | .051 | $F_{103,1}=20.8$ | **<10<sup>-3</sup>** | $F_{103,1}=30.9$ | **<10<sup>-3</sup>** | $F_{81,1}=5.0$ | **.028** | $F_{86,1}=0.6$ | .681 | $F_{106,1}=0.1$ | .985 | $F_{113,1}=0.9$ | .355 |
| $\bar{P}_{\text{M1L}}(\beta)$ | $F_{25,1}=2.0$ | .227 | $F_{102,1}=1.7$ | .245 | $F_{102,1}=0.0$ | .979 | $F_{26,1}=1.1$ | .412 | $F_{27,1}=0.9$ | .548 | $F_{102,1}=0.2$ | .820 | $F_{101,1}=1.0$ | .624 |
| $\bar{P}_{\text{M1R}}(\beta)$ | $F_{33,1}=2.0$ | .227 | $F_{102,1}=1.4$ | .245 | $F_{102,1}=0.2$ | .979 | $F_{28,1}=0.0$ | .998 | $F_{28,1}=0.7$ | .548 | $F_{102,1}=0.1$ | .820 | $F_{102,1}=0.2$ | .624 |
| $\bar{t}_{\text{M1L}}(\beta)$ | $F_{29,1}=8.8$ | **.010** | $F_{97,1}=15.2$ | **<10<sup>-3</sup>** | $F_{98,1}=24.9$ | **<10<sup>-3</sup>** | $F_{20,1}=9.6$ | **.008** | $F_{19,1}=0.1$ | .945 | $F_{103,1}=0.6$ | .889 | $F_{102,1}=0.3$ | .895 |
| $\bar{t}_{\text{M1R}}(\beta)$ | $F_{29,1}=9.0$ | **.010** | $F_{103,1}=18.7$ | **<10<sup>-3</sup>** | $F_{103,1}=19.6$ | **<10<sup>-3</sup>** | $F_{85,1}=11.2$ | **.005** | $F_{83,1}=0.2$ | .945 | $F_{104,1}=0.0$ | .968 | $F_{103,1}=0.1$ | .895 |
| $\bar{P}_{\text{PMC}_L}(\beta)$ | $F_{20,1}=2.2$ | .227 | $F_{101,1}=1.4$ | .245 | $F_{101,1}=0.1$ | .979 | $F_{23,1}=1.4$ | .412 | $F_{23,1}=0.9$ | .548 | $F_{101,1}=0.1$ | .820 | $F_{101,1}=0.4$ | .624 |
| $\bar{P}_{\text{PMC}_R}(\beta)$ | $F_{31,1}=0.2$ | .700 | $F_{102,1}=1.5$ | .245 | $F_{101,1}=0.0$ | .979 | $F_{34,1}=2.2$ | .412 | $F_{34,1}=0.0$ | .908 | $F_{102,1}=0.1$ | .820 | $F_{101,1}=0.6$ | .624 |
| $\bar{t}_{\text{PMC}_L}(\beta)$ | $F_{30,1}=7.6$ | **.010** | $F_{97,1}=16.4$ | **<10<sup>-3</sup>** | $F_{100,1}=25.2$ | **<10<sup>-3</sup>** | $F_{22,1}=9.2$ | **.008** | $F_{21,1}=0.0$ | .945 | $F_{97,1}=0.6$ | .889 | $F_{104,1}=0.2$ | .895 |
| $\bar{t}_{\text{PMC}_R}(\beta)$ | $F_{30,1}=7.5$ | **.010** | $F_{101,1}=18.3$ | **<10<sup>-3</sup>** | $F_{99,1}=22.9$ | **<10<sup>-3</sup>** | $F_{18,1}=4.3$ | .052 | $F_{18,1}=0.0$ | .945 | $F_{102,1}=0.2$ | .889 | $F_{100,1}=0.0$ | .895 |

\* LHDL = left-hand difficulty level, RHDL = right-hand difficulty level; all $p$ -values are FDR adjusted; bold-face indicates statistical significance.

The error of performance was significantly associated with the power modulation in bilateral M1 and in PMC_L_ in the beta frequency band, as well as in bilateral precuneus for the alpha frequency band, without any interactions between the power modulation, age, and/or task-related factors.

As illustrated in **Error! Reference source not found.** (bottom panel), the significant associations indicated a negative correlation between the performance error and the pseudo-*t*-values, implying that larger performance errors were associated with less normalized contrast between ERD-ERS modulations (Tables S.12 & S.13 in the *Supplementary Material*).

## Discussion

We investigated spectral differences in brain activity during unimanual and bimanual visuomotor tasks and their associations with the quality of behavioral performance in younger and older adults. We focused on the alpha and beta frequency bands at the source level, for which we estimated the mean power during dynamic and static pinch-grip force production and the corresponding (normalized) power modulation that we quantified through pseudo-*t*-values between these different phases of performance. The latter measure can be seen as a proxy for the size of modulation between event-related desynchronization and synchronization that occurred during dynamic and static motor activity, respectively. To disentangle age-specific power alterations, we also searched for possible associations between both spectral measures and the error of performance, which, by hypothesis, should be manifested in premotor areas.

As expected, our experimental protocol was accompanied by age-specific differences in the behavioral error scores. In the bimanual conditions, the total error (ℰ_Euclidean_) as well as the hand side-specific ones (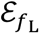 and 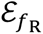) were highest during the performance of asymmetric motor tasks compared to symmetric ones. In these cases, the induced error on one side apparently spilled over to the other side, as both errors clearly increased even if we only increased the level of task difficulty for one side. Older participants generally performed worse, irrespective of the required motor task.

The beamformer-based localization revealed significant task-related sources for the beta frequency band in the motor network (bilateral primary and premotor areas) and for the alpha frequency band in bilateral (pre-)cuneus. In the latter, we identified significant age effects for the mean power and for the power modulations across performances, as both reduced with increasing age during unimanual and bimanual performance. By contrast, in the motor network, only the mean power in PMC_L_ significantly differed between age groups, and that only during bimanual performance; there again, the power was lower in older adults than in younger ones. LME models for the association between motor performance and spectral power signified a negative relationship between ℰ_Euclidean_ and the pseudo-*t*-values in both frequency bands over bilateral M1 and (pre-)cuneus and PMC_L_ during the bimanual tasks, as well as in bilateral M1 and precuneus_R_ during the unimanual tasks: Regardless of age, larger performance errors were associated with lower ERS/ERD power modulations.

### Motor performance

In line with previous studies, the quality of motor performance was significantly reduced with increasing age (<u>Babaeeghazvini et al., 2018</u>; <u>Chettouf et al., 2022</u>; <u>Lin et al., 2014</u>; <u>Rudisch et</u> <u>al., 2020</u>; <u>Swinnen, 1998</u>). Changes in left- and right-hand difficulty level significantly affected the error of performance. In the bimanual tasks, this was particularly pronounced during asymmetric tasks as shown by the significant interactions between left- and right-hand difficulty level in all performance measures, including the side-specific ones, 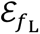 and 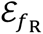. Increasing the difficulty level on one hand increased the error of performance not only at that side but also that of the other (Figure 10, middle and right panel; Table 5, 2^nd^ and 3^rd^ row). While other studies already reported asymmetric tasks to be more challenging (<u>Babaeeghazvini et al., 2018</u>; <u>Serrien and Wiesendanger, 2001</u>; <u>Solesio-Jofre et al., 2014</u>), our finding confirms a clear interference between the performance of the left and the right hand (<u>Cattaert et al., 1999</u>) in agreement with e.g., <u>Kennedy et al. (2016)</u>, who also found that the performance of one hand affects that of the other during asymmetric coordination tasks; see also (<u>Wenderoth et al., 2005</u>; <u>Wenderoth et al., 2004</u>; <u>Wenderoth et al., 2006</u>) [REFs]

### Alpha activity in visual areas

The time-frequency analysis over parietal and occipital electrodes unraveled clear changes in alpha power (8-14 Hz) between the epochs of static and dynamic force production. This was confirmed by the beamformers with which we could identify significant activity in occipital area, in (pre-)cuneus, and in (sensori)motor areas (Figure S.2). In the (pre-)cuneus, alpha activity differed significantly between age groups, with significantly lower alpha power during performance in older adults than in younger ones. This agrees with fMRI findings in which older adults showed elevated BOLD activity in the (pre-)cuneus during the exertion of isometric forces in unimanual precision grip tasks (<u>Godde et al., 2018</u>) or during simple motor tasks (<u>Hutchinson et al., 2002</u>; <u>Mattay et al., 2002</u>; <u>Ward and Frackowiak, 2003</u>) – recall that parietal and occipital alpha power is negatively correlated with BOLD during rest and motor activity alike (<u>Chettouf et al., 2022</u>; <u>de Munck et al., 2009</u>; <u>Gonçalves et al., 2003</u>).

Our LME models revealed not only a significant age effect on alpha mean power and power modulation in the bilateral (pre-)cuneus in unimanual and bimanual performance but also a strong association between the power modulation and the quality of performance (Tables 4 & 6). Across the board, the older adults displayed a lower ERS/ERD power modulation than the younger adults and performed worse. This suggests an altered visuospatial processing during the task performance (<u>Grefkes et al., 2004</u>; <u>Seitz and Binkofski, 2003</u>; <u>Zheng et al., 2023</u>). The (pre-)cuneus is commonly believed to be involved in the integration of visual and/or somatosensory information to guide motor actions (<u>Ashe and Georgopoulos, 1994</u>; <u>Cavada and</u> <u>Goldman-Rakic, 1989</u>; <u>Ferraina and Bianchi, 1994</u>; <u>Huttunen et al., 1996</u>; <u>Kalaska et al., 1990</u>; <u>Rizzolatti et al., 1998</u>; <u>Scott et al., 1997</u>). It also appears to be related to the attentional effort required for specific motor skills (<u>García-Larrea et al., 1991</u>; <u>Hari et al., 1990</u>; <u>Lam et al., 1999</u>; <u>Mima et al., 1998</u>). A reduced mean alpha power has been related to higher attention and engagement with the task (<u>Haegens et al., 2011</u>; <u>Rueda-Delgado et al., 2019</u>), as well as a decreased ability to switch between movement phases (<u>Deiber et al., 2012</u>; <u>Neubauer and Fink,</u> <u>2009</u>). Particularly interesting in this context is <u>Wenderoth et al. (2005)</u>’s conjecture that the (pre-)cuneus plays a role in the coordination effort required during bimanual movement and may contribute to a shift in attention between different locations in space, which is necessary to monitor the trajectories on the screen and may explain its activity during visuomotor tasks in our study. We can support this, given the larger errors of performance in the asymmetric visuomotor tasks that were accompanied by a significant reduction in alpha mean power in the (pre-)cuneus when compared to the symmetric visuomotor tasks.

<u>Margulies et al. (2009)</u> demonstrated that, next to being connected with adjacent visual cortical regions, the (pre-)cuneus displays other functional connectivity patterns. The central part is connected to the dorsolateral prefrontal, dorsomedial prefrontal, and multimodal lateral inferior parietal cortex, suggesting a cognitive/associative role. And, the anterior part of the (pre-)cuneus is connected with the superior parietal cortex, paracentral lobule, and motor cortex, suggesting a sensorimotor role – see also (<u>Cavanna and Trimble, 2006</u>; <u>Jitsuishi and Yamaguchi, 2023</u>; <u>Yamaguchi and Jitsuishi, 2024</u>). The here-observed, reduced alpha power modulation in older adults in the (pre-)cuneus could hence be interpreted as decreased responsiveness of the aged brain to visual stimuli during the visuomotor task (<u>Manor et al., 2023</u>). However, the reduced power modulation may also be a consequence of reduced capability to ‘switch’ from one task state to another. As such, it may be consistent with the reduced executive functions of older adults (including switching, inhibition, and working memory).

In any case, to align vision with motor performance, both sufficient excitation and inhibition in the (pre-)cuneus seem to be required (<u>Peng et al., 2015</u>). A significantly reduced alpha power modulation in this region may, hence, directly affect the quality of visuomotor performance in older adults. Interestingly, <u>Heuninckx et al. (2005)</u> reported higher precuneus BOLD activity in older than younger adults during (ipsilateral) coordination tasks, which might be a consequence of increased attentional deployment towards visuomotor task integration (consistent with our view). That is, the (pre-)cuneus might be a major hub region and a central core to manage this sensorimotor integration.

### Beta activity across the motor network

The 15-30 Hz beta power differed clearly between the dynamic and static force production phases, especially over central and frontocentral electrodes. The task-related beta activity was localized in bilateral SMA, M1, and PMC (e.g., <u>Gerloff and Andres, 2002</u>). However, unexpectedly, none of these areas showed any age-related differences during bimanual performance. The only exception was PMC_L_, which was much in line with our primary hypothesis. There, we found significantly lower beta mean power in older adults relative to younger ones. At first glance, this seemingly contradicts reports about increased beta power with age (<u>Karekal</u> <u>et al., 2023</u>), but this was observed during rest and interpreted as elevated cortical inhibition likely driven by GABAergic activity. The age-related reductions in motor-related mean beta power (<u>Chettouf et al., 2022</u>; <u>Park et al., 2024</u>; <u>Pascarella et al., 2025</u>) and post-movement (ERS) beta power (<u>Inamoto et al., 2023</u>; <u>Liu et al., 2017</u>) have been reported earlier and may be considered a sign of neurodegeneration in the aging brain (<u>Berger et al., 2020</u>; <u>Shih et al.,</u> <u>2021</u>). By the same token, however, a decreased mean beta power signifies elevated neural activity (<u>Pfurtscheller and Lopes da Silva, 1999</u>; <u>Rueda-Delgado et al., 2019</u>), suggesting altered inhibitory effects (increased excitation) (<u>Berger et al., 2020</u>; <u>Jensen and Mazaheri, 2010</u>) of PMC_L_ on ipsilateral M1 (<u>Daffertshofer et al., 2005</u>; <u>Stinear and Byblow, 2003</u>; <u>Stinear and</u> <u>Byblow, 2002</u>), or at least a compensatory effect of PMC. This result and interpretation align with previous studies that also found a significantly reduced mean beta power in PMC_L_ for older adults compared to younger adults during reaction time tasks (<u>Heinrichs-Graham and</u> <u>Wilson, 2016</u>), as well as during unimanual right-hand force grip tasks (<u>Berger et al., 2020</u>).

In PMC_L_ (and in bilateral M1), the ERD/ERS power modulations were significantly lower during asymmetric bimanual tasks compared to the symmetric ones. This finding also suggests that functional changes in PMC_L_ affect bilateral M1. While both the left and right PMCs are involved in bimanual coordination (<u>Beets et al., 2015</u>; <u>Debaere et al., 2004</u>; <u>Puttemans et al.,</u> <u>2005</u>), the PMC function is likely lateralized (<u>Beets et al., 2015</u>; <u>Debaere et al., 2004</u>; <u>Fujiyama</u> <u>et al., 2016a</u>; <u>Rueda-Delgado et al., 2017</u>; <u>Zivari Adab et al., 2018</u>). PMC_L_ plays a crucial role in bimanual performance accuracy by encoding the required speed of each hand (<u>Fujiyama et</u> <u>al., 2016a</u>), and in our study, PMC_L_ was significantly related to the quality of bimanual performance in both groups. Overall, our results support the involvement of the left hemisphere in bimanual performance as reported earlier (<u>Fujiyama et al., 2016a</u>; <u>Jäncke et al., 2000</u>; <u>Toyo-</u> <u>kura et al., 1999</u>) and, more specifically, PMC_L_. It seems that the left hemisphere contributes asymmetrically to bimanual coordination (<u>Merrick et al., 2022</u>) and more generally to left and right interlimb coordination (<u>Swinnen et al., 2010</u>).

As mentioned in the *Results* section regarding Figure 12, the reduction in mean power and power modulation in the left hemisphere of older adults appears to be accompanied by a corresponding increase in the right hemisphere. At first glance, one might consider this a compensatory activity, especially since the age differences in the left hemisphere appear to be just inverted from the right one. By the same token, the difference between left and right, i.e., lateralization, seems to be much reduced in the older age group. This observation may ‘explain’ earlier reports about a possible loss of hand dominance with advancing age (<u>Kalisch et al.,</u> <u>2006</u>), the preferred use of the dominant hand (in our case, primarily the right hand) alters towards a more ambidextrous use with increasing age. However, the differences that we observed in the right hemisphere did not reach significance, which led us to abstain from further speculation about this. Of note, however, a power decrease in one hemisphere and an increase in the other may contribute to some inconsistencies in reports about the mean beta power in M1 and PMC, as well as concurrent interpretations of their functional interaction and general role in coordinating the left and right hands.

#### Limitations

We applied linear mixed-effects models to investigate whether alpha and beta activities and their interactions with *age* and task-related factors predict performance error during unimanual and bimanual tasks. Overall, the reduced alpha and beta power modulations were significantly associated with larger performance errors in both groups. It seems that a sufficient difference between ERS and ERD power (pseudo-*t*-values) is needed for an appropriate quality of motor performance. However, we did not observe any task- or age-specific associations. This might be influenced by the fact that we selected significant clusters in each ROI for both age groups collectively, rather than specifying them for each age group individually.

The long duration of the experiment rendered the definition of a proper baseline a challenge. For the between-subject comparisons of the mean power, we opted for normalizing the EEG data by the mean standard deviation over all conditions, channels, and time points. By quantifying power modulations via the normalized difference between ERS- and ERD-related phases, the pseudo-*t*-value, we circumvented the issue altogether. However, this comes at the price of being unable to distinguish between the isolated effects of event-related desynchronization (ERD) vis-à-vis event-related synchronization (ERS).

## Conclusions

Bimanual coordination declined in older adults, especially when visuomotor task requirements were difficult and asymmetric. The differences in motor performance were accompanied by significant differences in both the mean power and power modulations in the alpha band in (pre-)cuneus, where both hallmark power values were significantly lower in older adults than in younger ones. The age-related differences in alpha power in (pre-)cuneus areas hint at a decreased responsiveness to visual stimuli during visuomotor tasks maybe due to the difficulty of older adults in suppressing unwanted information processing during motor performance. In the motor network we observed age-related decreases in mean beta power especially in PMC_L_. This agrees with an improperly timed inhibition of PMC_L_ on ipsilateral M1, which contributes to a reduced quality of motor performance in older adults. During unimanual performances, the ERD/ERS power modulations in bilateral M1 and (pre-)cuneus_R_ were negatively associated with the error of performance, and during bimanual performance we observed this also in PMC_L_ and (pre-)cuneus_L_. We can confirm the hypothesized necessity of appropriate ERD/ERS power modulation in the quality of motor performance and the pivotal role of PMC_L_ for bimanual performance. Age-related differences in (pre-)cuneus and PMC_L_ may jeopardize attentional deployment for visual processing, visual and motor network integration, and (bimanual) motor coordination alike.

## Appendix

### A – Illustration of the left/right force errors

**Figure A.1.**
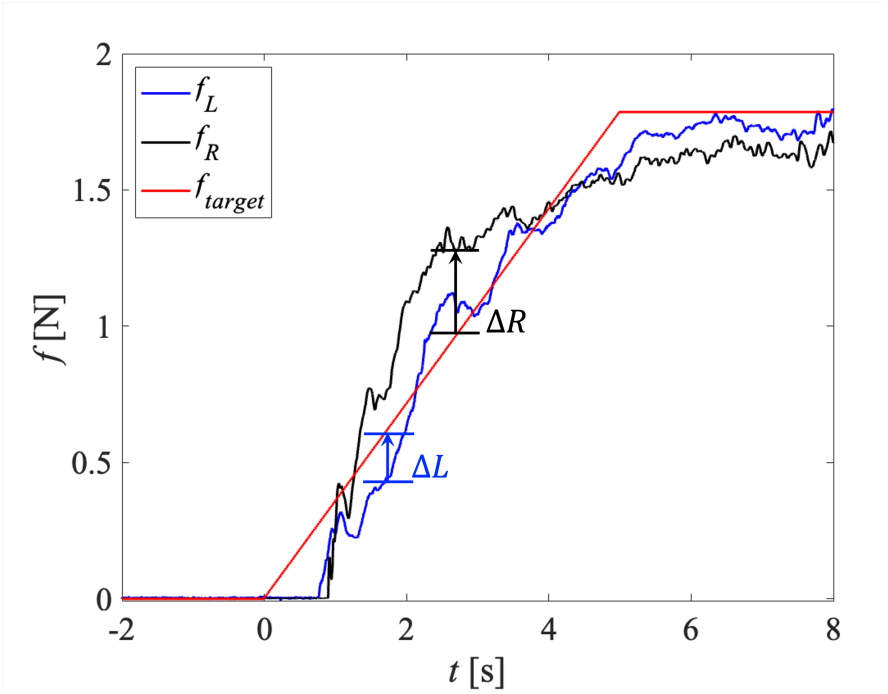
Like Figure 2 in the main text, we here illustrate the force difference of each hand with target force. Δ_*t*_is the difference between *f_L_* and *f_T_*. Δ_*R*_is the difference between *f_R_* and *f_T_*. Left force (*f_L_*), right force (*f_R_*), target force (*f_T_*).

### B – Source reconstruction and MNI coordinates

**Figure B.1.**
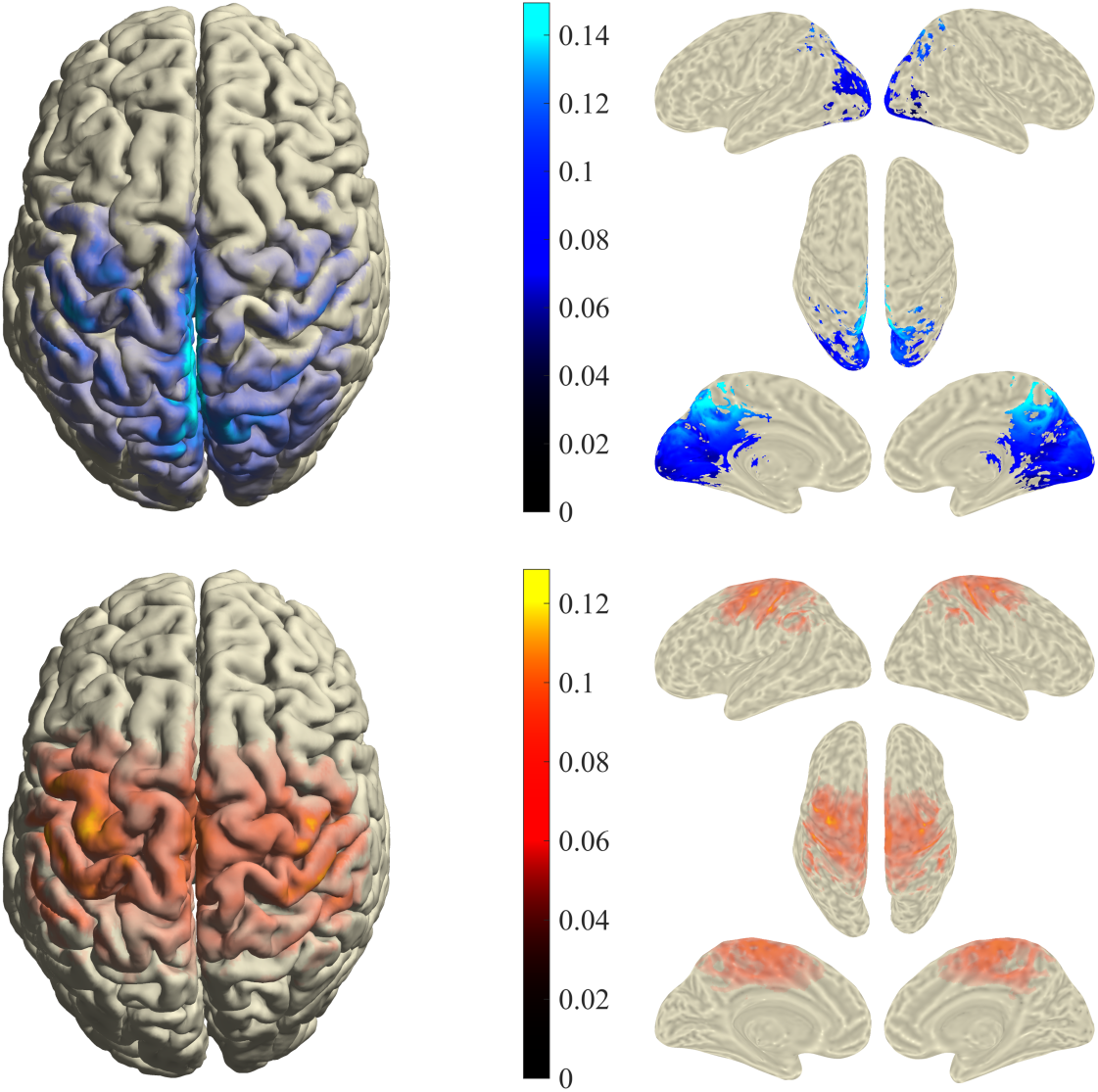
Power values of the beam-former statistics. In contrast to Figure 5 of the main text here we did not apply a mask based on significance. Opacity is set via the power values.

**Figure B.2.**
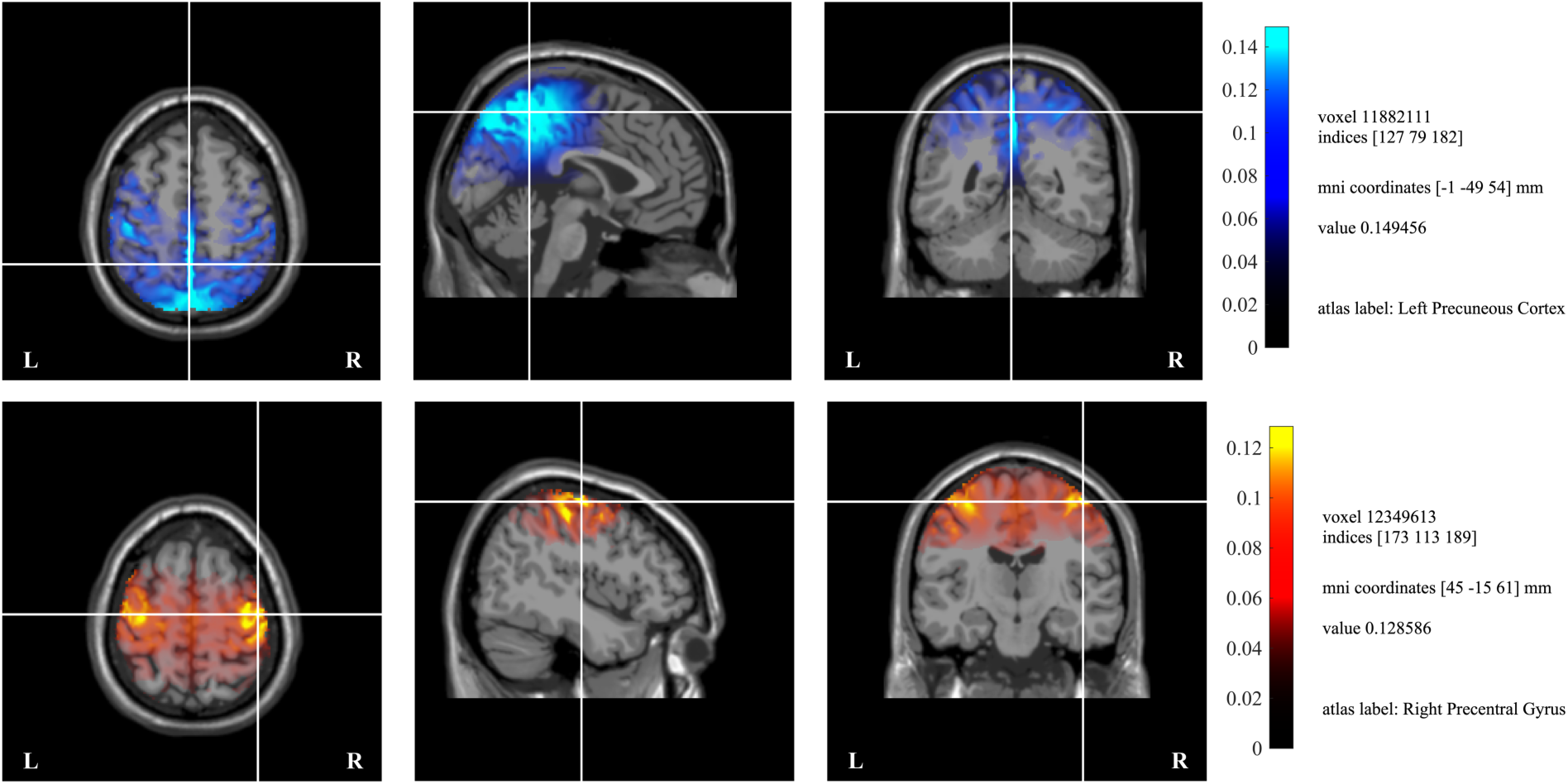
MNI coordination related to the maximum power values in the alpha band (upper row) and in the beta band (lower row); cf. Figure B.1.

**Figure B.3.**
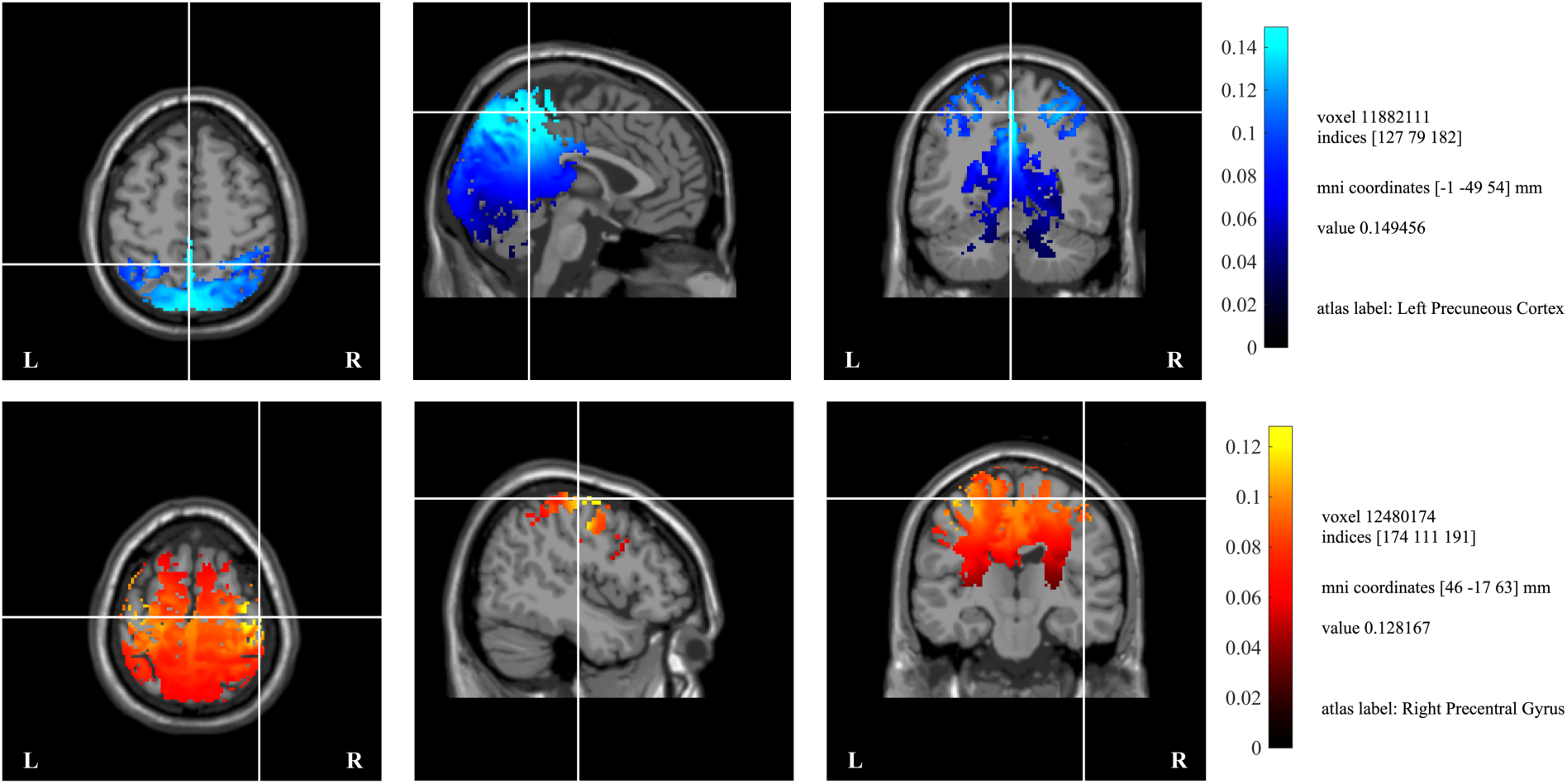
MNI coordinates related to the maximum significant *t*-value when estimating the effect of group in the alpha band (top row) and the effect of task in the beta band (bottom row); cf. Figure 5 in the main text.

