## Supplementary Material for "Age-related differences in the cortical motor network during unimanual and bimanual coordination – an EEG study"

### Trial characteristics

**Table S.1.** The number of trials per subject and the number of ICA components are considered artifacts and removed. Participants Y18, O09, and O17 were removed from further analysis (see the Error! **Reference source not found.** for exclusion criteria).

| ID | # trial per task/condition | | | | | | | | average # ICA artifacts | | |
|---|---|---|---|---|---|---|---|---|---|---|---|
| | *RN* | *RD* | *LN* | *LD* | *SN* | *SD* | *ASRD* | *ASLD* | *head* | *muscle* | *eye* |
| **Y01** | 20 | 20 | 20 | 20 | 30 | 30 | 30 | 30 | 1.00 | 4.00 | 5.00 |
| **Y02** | 20 | 20 | 20 | 20 | 30 | 29 | 30 | 30 | 1.00 | 0.00 | 5.33 |
| **Y03** | 20 | 18 | 18 | 20 | 30 | 28 | 30 | 29 | 1.00 | 0.00 | 5.33 |
| **Y04** | 20 | 20 | 20 | 20 | 30 | 30 | 30 | 30 | 0.00 | 0.67 | 6.67 |
| **Y05** | 20 | 20 | 20 | 19 | 30 | 25 | 30 | 28 | 0.33 | 2.67 | 6.00 |
| **Y06** | 18 | 13 | 14 | 20 | 22 | 24 | 15 | 21 | 1.00 | 7.67 | 3.67 |
| **Y07** | 20 | 20 | 20 | 19 | 30 | 30 | 30 | 30 | 1.00 | 0.00 | 3.67 |
| **Y08** | 20 | 20 | 20 | 20 | 30 | 30 | 28 | 29 | 0.00 | 0.00 | 5.67 |
| **Y09** | 19 | 20 | 20 | 20 | 30 | 30 | 30 | 30 | 0.33 | 0.00 | 4.33 |
| **Y10** | 20 | 20 | 20 | 20 | 30 | 30 | 30 | 30 | 1.00 | 0.33 | 4.67 |
| **Y11** | 20 | 18 | 20 | 20 | 30 | 26 | 29 | 25 | 0.00 | 0.33 | 8.33 |
| **Y12** | 14 | 11 | 13 | 17 | 16 | 17 | 12 | 17 | 0.33 | 9.67 | 4.33 |
| **Y13** | 30 | 29 | 30 | 30 | 30 | 30 | 30 | 30 | 0.00 | 1.67 | 6.67 |
| **Y14** | 29 | 30 | 30 | 30 | 27 | 29 | 30 | 29 | 0.00 | 3.67 | 7.67 |
| **Y15** | 26 | 27 | 27 | 26 | 26 | 24 | 26 | 30 | 0.00 | 0.00 | 3.33 |
| **Y16** | 30 | 30 | 30 | 30 | 30 | 30 | 30 | 30 | 1.00 | 0.33 | 4.67 |
| ~~**Y17**~~ | ~~30~~ | ~~30~~ | ~~29~~ | ~~30~~ | ~~30~~ | ~~30~~ | ~~30~~ | ~~30~~ | ~~0.00~~ | ~~1.00~~ | ~~4.33~~ |
| **Y18** | 0 | 10 | 7 | 19 | ~~19~~ | ~~10~~ | ~~10~~ | ~~19~~ | ~~1.00~~ | ~~0.00~~ | ~~5.00~~ |
| **Y19** | 28 | 28 | 27 | 29 | 25 | 27 | 29 | 27 | 0.00 | 1.00 | 3.33 |
| **O01** | 20 | 20 | 20 | 20 | 30 | 30 | 30 | 30 | 1.00 | 0.00 | 5.67 |
| **O02** | 20 | 20 | 20 | 20 | 30 | 30 | 30 | 30 | 0.00 | 14.67 | 2.67 |
| **O03** | 20 | 10 | 9 | 10 | 20 | 20 | 19 | 20 | 0.00 | 1.00 | 3.50 |
| **O04** | 7 | 20 | 10 | 20 | 19 | 12 | 10 | 15 | 1.00 | 6.00 | 4.00 |
| **O05** | 9 | 29 | 18 | 19 | 30 | 28 | 26 | 29 | 0.00 | 0.67 | 3.33 |
| **O06** | 10 | 20 | 10 | 20 | 20 | 20 | 20 | 20 | 0.00 | 11.50 | 5.50 |
| **O07** | 10 | 30 | 19 | 20 | 30 | 30 | 29 | 30 | 0.00 | 4.33 | 6.00 |
| ~~**O08**~~ | ~~9~~ | ~~30~~ | ~~20~~ | ~~20~~ | ~~30~~ | ~~30~~ | ~~26~~ | ~~30~~ | ~~0.00~~ | ~~1.33~~ | ~~4.00~~ |
| **O09** | 10 | 29 | 20 | 19 | 28 | ~~28~~ | ~~26~~ | ~~28~~ | ~~0.00~~ | ~~6.00~~ | ~~4.00~~ |
| **O10** | 10 | 30 | 19 | 20 | 30 | 29 | 30 | 30 | 0.00 | 0.00 | 4.00 |
| **O11** | 10 | 18 | 10 | 9 | 19 | 15 | 12 | 12 | 0.00 | 1.50 | 6.50 |
| **O12** | 29 | 26 | 27 | 28 | 29 | 30 | 29 | 21 | 0.00 | 0.33 | 6.00 |
| **O13** | 30 | 30 | 30 | 30 | 30 | 30 | 30 | 30 | 1.00 | 1.00 | 3.67 |
| **O14** | 25 | 30 | 26 | 25 | 27 | 23 | 24 | 27 | 0.67 | 2.67 | 3.67 |
| **O15** | 30 | 30 | 29 | 29 | 30 | 29 | 29 | 30 | 0.00 | 3.33 | 4.67 |
| **O16** | 25 | 28 | 20 | 25 | 28 | 25 | 19 | 25 | 0.33 | 12.00 | 4.00 |
| ~~**O17**~~ | ~~27~~ | ~~30~~ | ~~30~~ | ~~30~~ | ~~27~~ | ~~26~~ | ~~2~~ | ~~16~~ | ~~1.00~~ | ~~2.33~~ | ~~6.67~~ |
| **O18** | 29 | 19 | 30 | 30 | 29 | 30 | 29 | 30 | 0.00 | 0.00 | 3.33 |
| **O19** | 30 | 30 | 27 | 29 | 30 | 28 | 29 | 28 | 0.00 | 1.33 | 3.67 |

### Behavioral characteristics

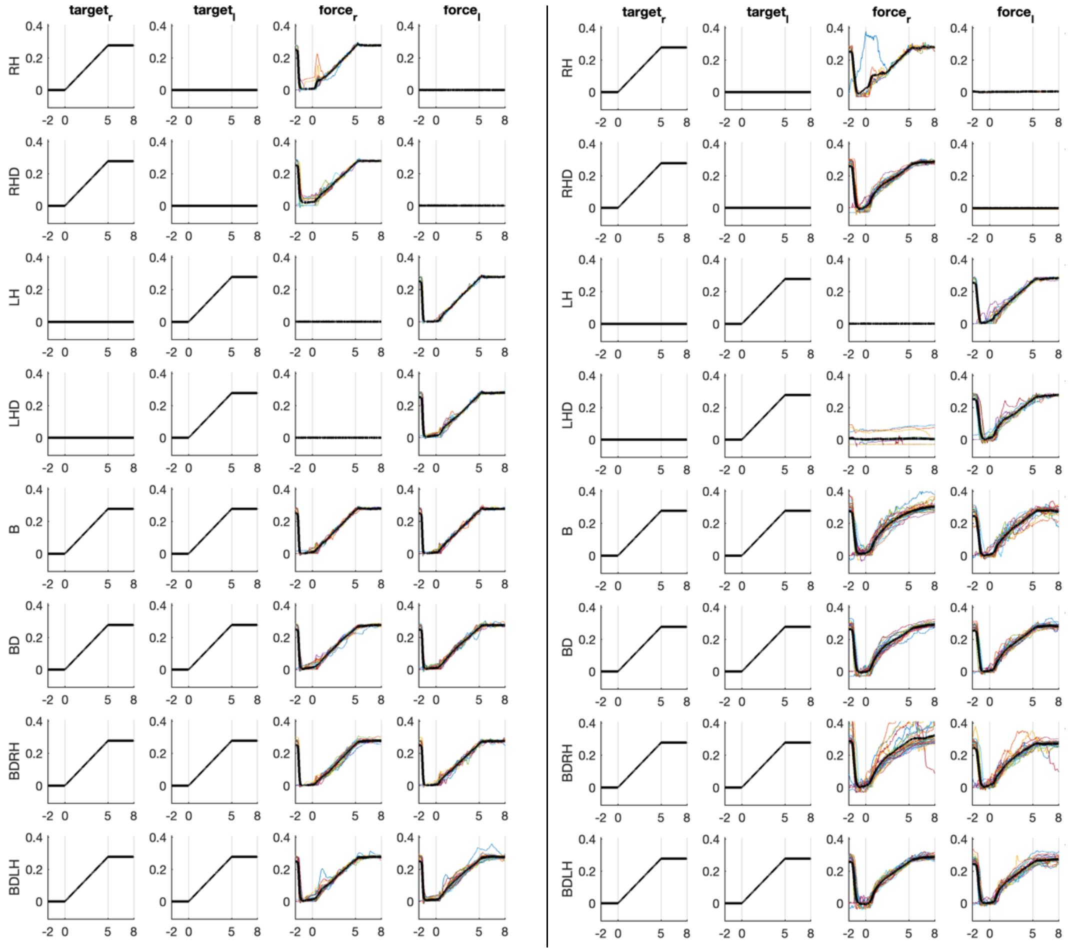


**Figure S.1**. Two examples of trial performance in different tasks. The figure shows the level of force applied on the sensors in two random participants from young (left panel) and older adults (right panel).

### Alpha activity – beamformer statistics

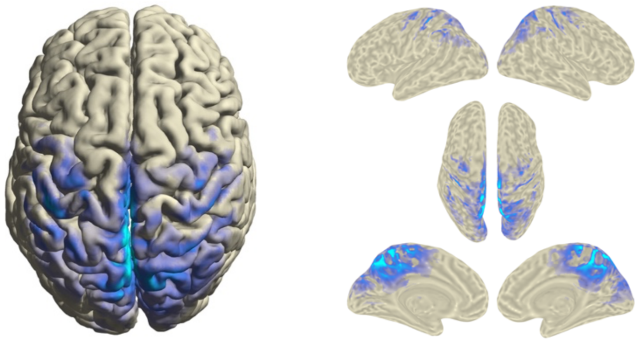

**Figure S.2**. Alpha activity projected to the cortical surface. The color-coding is based on the resulting *t*-statistic for the main effect of task. Like Figure 4, both bilateral precuneus and bilateral cuneal cortex show strong contrasts between the static and the dynamic phases; in Figure A.3 we also show selected MRI slices including the MNI coordinates.

### Statistics of the unimanual tasks*

**Table S.2.** Main effects of the mean alpha power and pseudo-*t*-values.

| contrast | estimate | SE | df | *t*-value | *p*-value |
|---|---|---|---|---|---|
| $\bar{\boldsymbol{P}}_{\mathbf{prec}_L}(\alpha)$ younger – older | 6.12 | 1.36 | 33.00 | 4.48 | $<10^{-3}$ |
| $\bar{\boldsymbol{P}}_{\mathbf{prec}_L}(\alpha)$ normal – difficult (HDL) | 0.23 | 0.09 | 99.00 | 2.50 | 0.01 |
| $\bar{\boldsymbol{P}}_{\mathbf{prec}_R}(\alpha)$ younger – older | 5.54 | 1.45 | 33.00 | 3.82 | $<10^{-3}$ |
| $\bar{\boldsymbol{P}}_{\mathbf{prec}_R}(\alpha)$ normal – difficult (HDL) | 0.29 | 0.11 | 99.00 | 2.66 | 0.01 |
| $\bar{\boldsymbol{t}}_{\mathbf{prec}_L}(\alpha)$ younger – older | 0.09 | 0.02 | 33.00 | 4.07 | $<10^{-3}$ |
| $\bar{\boldsymbol{t}}_{\mathbf{prec}_L}(\alpha)$ normal – difficult (HDL) | 0.04 | 0.01 | 99.00 | 3.49 | $<10^{-3}$ |
| $\bar{\boldsymbol{t}}_{\mathbf{prec}_R}(\alpha)$ younger – older | 0.09 | 0.02 | 33.00 | 3.87 | $<10^{-3}$ |
| $\bar{\boldsymbol{t}}_{\mathbf{prec}_R}(\alpha)$ normal – difficult (HDL) | 0.04 | 0.01 | 99.00 | 2.82 | 0.01 |

**Table S.3.** Interaction effects of the mean alpha power.

| contrast | HS/age | estimate | SE | df | *t*-value | *p*-value |
|---|---|---|---|---|---|---|
| $\bar{\boldsymbol{P}}_{\mathbf{prec}_L}(\alpha)$ younger - older | right | 6.31 | 1.37 | 33.29 | 4.61 | $<10^{-3}$ |
| $\bar{\boldsymbol{P}}_{\mathbf{prec}_L}(\alpha)$ younger - older | left | 5.93 | 1.37 | 33.29 | 4.34 | $<10^{-3}$ |
| $\bar{\boldsymbol{P}}_{\mathbf{prec}_R}(\alpha)$ younger – older | right | 5.80 | 1.45 | 33.38 | 3.99 | $<10^{-3}$ |
| $\bar{\boldsymbol{P}}_{\mathbf{prec}_R}(\alpha)$ younger – older | left | 5.27 | 1.45 | 33.38 | 3.63 | $<10^{-3}$ |
| $\bar{\boldsymbol{P}}_{\mathbf{prec}_L}(\alpha)$ right – left (HS) | younger | 0.25 | 0.13 | 99.00 | 1.99 | 0.05 |
| $\bar{\boldsymbol{P}}_{\mathbf{prec}_L}(\alpha)$ right – left (HS) | older | -0.13 | 0.13 | 99.00 | -0.99 | 0.32 |
| $\bar{\boldsymbol{P}}_{\mathbf{prec}_R}(\alpha)$ right – left (HS) | younger | 0.45 | 0.15 | 99.00 | 2.92 | $<10^{-3}$ |
| $\bar{\boldsymbol{P}}_{\mathbf{prec}_R}(\alpha)$ right – left (HS) | older | -0.08 | 0.16 | 99.00 | -0.52 | 0.61 |
| $\bar{\boldsymbol{P}}_{\mathbf{prec}_L}(\alpha)$ normal – difficult (HDL) | right | -0.01 | 0.13 | 99.00 | -0.06 | 0.95 |
| $\bar{\boldsymbol{P}}_{\mathbf{prec}_L}(\alpha)$ normal – difficult (HDL) | left | 0.46 | 0.13 | 99.00 | 3.60 | $<10^{-3}$ |
| $\bar{\boldsymbol{P}}_{\mathbf{prec}_L}(\alpha)$ right – left (HS) | normal | -0.17 | 0.13 | 99.00 | -1.35 | 0.18 |
| $\bar{\boldsymbol{P}}_{\mathbf{prec}_L}(\alpha)$ right – left (HS) | difficult | 0.30 | 0.13 | 99.00 | 2.31 | 0.02 |

**Table S.4.** Main effects of mean beta power and pseudo-*t*-values.

| contrast | estimate | SE | df | *t*-value | *p*-value |
|---|---|---|---|---|---|
| $\bar{\boldsymbol{t}}_{\boldsymbol{M1}_L}(\boldsymbol{\beta})$ normal – difficult (HDL) | 0.04 | 0.01 | 99.00 | 3.32 | $<10^{-3}$ |
| $\bar{\boldsymbol{t}}_{\boldsymbol{M1}_R}(\boldsymbol{\beta})$ normal – difficult (HDL) | 0.04 | 0.01 | 99.00 | 3.11 | $<10^{-3}$ |
| $\bar{\boldsymbol{t}}_{\boldsymbol{PMC}_L}(\boldsymbol{\beta})$ normal – difficult (HDL) | 0.03 | 0.01 | 99.00 | 2.78 | 0.01 |
| $\bar{\boldsymbol{t}}_{\boldsymbol{PMC}_R}(\boldsymbol{\beta})$ normal – difficult (HDL) | 0.02 | 0.01 | 99.00 | 2.07 | 0.04 |
| $\bar{\boldsymbol{P}}_{\boldsymbol{M1}_L}(\boldsymbol{\beta})$ right – left (HS) | -0.23 | 0.08 | 99.00 | -2.92 | $<10^{-3}$ |
| $\bar{\boldsymbol{P}}_{\boldsymbol{PMC}_L}(\boldsymbol{\beta})$ right – left (HS) | -0.23 | 0.09 | 99.00 | -2.66 | 0.01 |
| $\bar{\boldsymbol{t}}_{\boldsymbol{M1}_R}(\boldsymbol{\beta})$ right – left (HS) | -0.06 | 0.01 | 99.00 | -4.36 | $<10^{-3}$ |
| $\bar{\boldsymbol{t}}_{\boldsymbol{PMC}_R}(\boldsymbol{\beta})$ right – left (HS) | -0.04 | 0.01 | 99.00 | -3.24 | $<10^{-3}$ |

**Table S.5.** Interaction effects of the mean beta power and pseudo-*t*-value.

| contrast | HS/age | estimate | SE | df | *t*-value | *p*-value |
|---|---|---|---|---|---|---|
| $\bar{\boldsymbol{P}}_{\boldsymbol{M1}_L}(\boldsymbol{\beta})$ younger - older | right | 2.67 | 1.10 | 33.33 | 2.43 | 0.02 |
| $\bar{\boldsymbol{P}}_{\boldsymbol{M1}_L}(\boldsymbol{\beta})$ younger - older | left | 2.28 | 1.10 | 33.33 | 2.07 | 0.05 |
| $\bar{\boldsymbol{P}}_{\boldsymbol{M1}_R}(\boldsymbol{\beta})$ right – left (HS) | younger | 0.42 | 0.13 | 99.00 | 3.24 | $<10^{-3}$ |
| $\bar{\boldsymbol{P}}_{\boldsymbol{M1}_R}(\boldsymbol{\beta})$ right – left (HS) | older | -0.06 | 0.13 | 99.00 | -0.49 | 0.63 |
| $\bar{\boldsymbol{P}}_{\boldsymbol{PMC}_L}(\boldsymbol{\beta})$ younger - older | right | 3.30 | 1.18 | 33.36 | 2.80 | 0.01 |
| $\bar{\boldsymbol{P}}_{\boldsymbol{PMC}_L}(\boldsymbol{\beta})$ younger - older | left | 2.84 | 1.18 | 33.36 | 2.41 | 0.02 |
| $\bar{\boldsymbol{P}}_{\boldsymbol{PMC}_R}(\boldsymbol{\beta})$ right – left (HS) | younger | 0.33 | 0.12 | 99.00 | 2.77 | 0.01 |
| $\bar{\boldsymbol{P}}_{\boldsymbol{PMC}_R}(\boldsymbol{\beta})$ right – left (HS) | older | -0.16 | 0.12 | 99.00 | -1.27 | 0.21 |
| $\bar{\boldsymbol{t}}_{\boldsymbol{M1}_R}(\boldsymbol{\beta})$ right – left (HS) | younger | -0.05 | 0.02 | 99.00 | -2.50 | 0.01 |
| $\bar{\boldsymbol{t}}_{\boldsymbol{M1}_R}(\boldsymbol{\beta})$ right – left (HS) | older | -0.07 | 0.02 | 99.00 | -3.66 | $<10^{-3}$ |
| $\bar{\boldsymbol{t}}_{\boldsymbol{PMC}_R}(\boldsymbol{\beta})$ right – left (HS) | younger | -0.02 | 0.02 | 99.00 | -1.34 | 0.18 |
| $\bar{\boldsymbol{t}}_{\boldsymbol{PMC}_R}(\boldsymbol{\beta})$ right – left (HS) | older | -0.06 | 0.02 | 99.00 | -3.21 | $<10^{-3}$ |
| $\bar{\boldsymbol{P}}_{\boldsymbol{M1}_L}(\boldsymbol{\beta})$ right – left (HS) | normal | -0.47 | 0.11 | 99.00 | -4.27 | $<10^{-3}$ |
| $\bar{\boldsymbol{P}}_{\boldsymbol{M1}_L}(\boldsymbol{\beta})$ right – left (HS) | difficult | 0.02 | 0.11 | 99.00 | 0.14 | 0.89 |
| $\bar{\boldsymbol{P}}_{\boldsymbol{M1}_R}(\boldsymbol{\beta})$ right – left (HS) | normal | -0.07 | 0.13 | 99.00 | -0.53 | 0.60 |
| $\bar{\boldsymbol{P}}_{\boldsymbol{M1}_R}(\boldsymbol{\beta})$ right – left (HS) | difficult | 0.42 | 0.13 | 99.00 | 3.23 | $<10^{-3}$ |
| $\bar{\boldsymbol{P}}_{\boldsymbol{PMC}_L}(\boldsymbol{\beta})$ right – left (HS) | normal | -0.52 | 0.12 | 99.00 | -4.26 | $<10^{-3}$ |
| $\bar{\boldsymbol{P}}_{\boldsymbol{PMC}_L}(\boldsymbol{\beta})$ right – left (HS) | difficult | 0.06 | 0.12 | 99.00 | 0.50 | 0.62 |
| $\bar{\boldsymbol{P}}_{\boldsymbol{PMC}_R}(\boldsymbol{\beta})$ right – left (HS) | normal | -0.16 | 0.12 | 99.00 | -1.33 | 0.19 |
| $\bar{\boldsymbol{P}}_{\boldsymbol{PMC}_R}(\boldsymbol{\beta})$ right – left (HS) | difficult | 0.34 | 0.12 | 99.00 | 2.77 | 0.01 |

* Abbreviations: Standard error (SE); degree of freedom (df); hand difficulty level (HDL); hand side (HS).

| | | | | | | |
|---|---|---|---|---|---|---|
| $\bar{\boldsymbol{t}}_{\boldsymbol{M1_L}}(\boldsymbol{\beta})$ right – left (HS) | normal | -0.05 | 0.02 | 99.00 | -2.74 | 0.01 |
| $\bar{\boldsymbol{t}}_{\boldsymbol{M1_L}}(\boldsymbol{\beta})$ right – left (HS) | difficult | 0.03 | 0.02 | 99.00 | 1.55 | 0.12 |
| $\bar{\boldsymbol{t}}_{\boldsymbol{PMC_L}}(\boldsymbol{\beta})$ right – left (HS) | normal | -0.04 | 0.02 | 99.00 | -2.49 | 0.01 |
| $\bar{\boldsymbol{t}}_{\boldsymbol{PMC_L}}(\boldsymbol{\beta})$ right – left (HS) | difficult | 0.03 | 0.02 | 99.00 | 1.60 | 0.11 |

### Statistics of the bimanual tasks[†]

**Table S.6.** Main effects of $\boldsymbol{\mathcal{E}}_{\mathbf{Euclidean}}$, $\boldsymbol{\mathcal{E}}_{\boldsymbol{f_L}}$ and $\boldsymbol{\mathcal{E}}_{\boldsymbol{f_R}}$.

| contrast | estimate | SE | df | *t*-value | *p*-value |
|---|---|---|---|---|---|
| $\boldsymbol{\mathcal{E}}_{\mathbf{Euclidean}}$ younger - older | -0.53 | 0.13 | 33.00 | -4.15 | $<10^{-3}$ |
| $\boldsymbol{\mathcal{E}}_{\mathbf{Euclidean}}$ normal – difficult (LHDL) | -0.40 | 0.03 | 99.00 | -15.19 | $<10^{-3}$ |
| $\boldsymbol{\mathcal{E}}_{\mathbf{Euclidean}}$ normal – difficult (RHDL) | -0.43 | 0.03 | 99.00 | -16.40 | $<10^{-3}$ |
| $\boldsymbol{\mathcal{E}}_{\boldsymbol{f_L}}$ younger - older | -0.43 | 0.13 | 33.00 | -3.44 | $<10^{-3}$ |
| $\boldsymbol{\mathcal{E}}_{\boldsymbol{f_L}}$ normal – difficult (LHDL) | -0.36 | 0.03 | 99.00 | -12.33 | $<10^{-3}$ |
| $\boldsymbol{\mathcal{E}}_{\boldsymbol{f_L}}$ normal – difficult (RHDL) | 0.15 | 0.03 | 99.00 | 5.12 | $<10^{-3}$ |
| $\boldsymbol{\mathcal{E}}_{\boldsymbol{f_R}}$ younger - older | -0.45 | 0.12 | 33.00 | -3.62 | $<10^{-3}$ |
| $\boldsymbol{\mathcal{E}}_{\boldsymbol{f_R}}$ normal – difficult (LHDL) | 0.16 | 0.03 | 99.00 | 5.90 | $<10^{-3}$ |
| $\boldsymbol{\mathcal{E}}_{\boldsymbol{f_R}}$ normal – difficult (RHDL) | -0.42 | 0.03 | 99.00 | -15.65 | $<10^{-3}$ |

**Table S.7.** Interaction effects of $\boldsymbol{\mathcal{E}}_{\mathbf{Euclidean}}$, $\boldsymbol{\mathcal{E}}_{\boldsymbol{f_L}}$ and $\boldsymbol{\mathcal{E}}_{\boldsymbol{f_R}}$.

| contrast | LHDL/RHDL | estimate | SE | df | t-value | p-value |
|---|---|---|---|---|---|---|
| $\boldsymbol{\mathcal{E}}_{\mathbf{Euclidean}}$ normal – difficult (RHDL) | normal | -1.03 | 0.04 | 99.00 | -27.76 | $<10^{-3}$ |
| $\boldsymbol{\mathcal{E}}_{\mathbf{Euclidean}}$ normal – difficult (RHDL) | difficult | 0.17 | 0.04 | 99.00 | 4.56 | $<10^{-3}$ |
| $\boldsymbol{\mathcal{E}}_{\mathbf{Euclidean}}$ normal – difficult (LHDL) | normal | -1.00 | 0.04 | 99.00 | -26.90 | $<10^{-3}$ |
| $\boldsymbol{\mathcal{E}}_{\mathbf{Euclidean}}$ normal – difficult (LHDL) | difficult | 0.20 | 0.04 | 99.00 | 5.42 | $<10^{-3}$ |
| $\boldsymbol{\mathcal{E}}_{\boldsymbol{f_L}}$ normal – difficult (RHDL) | normal | -0.20 | 0.04 | 99.00 | -4.91 | $<10^{-3}$ |
| $\boldsymbol{\mathcal{E}}_{\boldsymbol{f_L}}$ normal – difficult (RHDL) | difficult | 0.51 | 0.04 | 99.00 | 12.15 | $<10^{-3}$ |
| $\boldsymbol{\mathcal{E}}_{\boldsymbol{f_R}}$ normal – difficult (LHDL) | normal | -0.20 | 0.04 | 99.00 | -5.25 | $<10^{-3}$ |
| $\boldsymbol{\mathcal{E}}_{\boldsymbol{f_R}}$ normal – difficult (LHDL) | difficult | 0.52 | 0.04 | 99.00 | 13.60 | $<10^{-3}$ |

**Table S.8.** Main effects of the mean alpha power and pseudo-*t*-value.

| contrast | estimate | SE | df | *t*-value | *p*-value |
|---|---|---|---|---|---|
| $\bar{\boldsymbol{P}}_{\mathbf{prec_L}}(\alpha)$ younger – older | 5.79 | 1.35 | 33.00 | 4.29 | $<10^{-3}$ |
| $\bar{\boldsymbol{P}}_{\mathbf{prec_R}}(\alpha)$ younger – older | 5.19 | 1.46 | 33.00 | 3.57 | $<10^{-3}$ |
| $\bar{\boldsymbol{t}}_{\mathbf{prec_L}}(\alpha)$ younger – older | 0.12 | 0.03 | 33.00 | 4.39 | $<10^{-3}$ |
| $\bar{\boldsymbol{t}}_{\mathbf{prec_R}}(\alpha)$ younger – older | 0.12 | 0.03 | 33.00 | 4.76 | $<10^{-3}$ |

**Table S.9.** Interaction effects of the mean alpha power and pseudo-*t*-value.

| contrast | LHDL | estimate | SE | df | *t*-value | *p*-value |
|---|---|---|---|---|---|---|
| $\bar{\boldsymbol{P}}_{\mathbf{prec_L}}(\alpha)$ normal – difficult (RHDL) | normal | 0.27 | 0.08 | 99.00 | 3.31 | $<10^{-3}$ |
| $\bar{\boldsymbol{P}}_{\mathbf{prec_L}}(\alpha)$ normal – difficult (RHDL) | difficult | -0.06 | 0.08 | 99.00 | -0.75 | 0.46 |
| $\bar{\boldsymbol{P}}_{\mathbf{prec_R}}(\alpha)$ normal – difficult (RHDL) | normal | 0.26 | 0.09 | 99.00 | 2.94 | $<10^{-3}$ |
| $\bar{\boldsymbol{P}}_{\mathbf{prec_R}}(\alpha)$ normal – difficult (RHDL) | difficult | -0.01 | 0.09 | 99.00 | -0.17 | 0.87 |
| $\bar{\boldsymbol{t}}_{\mathbf{prec_L}}(\alpha)$ normal – difficult (RHDL) | normal | 0.05 | 0.02 | 99.00 | 3.07 | $<10^{-3}$ |
| $\bar{\boldsymbol{t}}_{\mathbf{prec_L}}(\alpha)$ normal – difficult (RHDL) | difficult | -0.01 | 0.02 | 99.00 | -0.74 | 0.46 |
| $\bar{\boldsymbol{t}}_{\mathbf{prec_R}}(\alpha)$ normal – difficult (RHDL) | normal | 0.06 | 0.02 | 99.00 | 3.34 | $<10^{-3}$ |
| $\bar{\boldsymbol{t}}_{\mathbf{prec_R}}(\alpha)$ normal – difficult (RHDL) | difficult | 0.00 | 0.02 | 99.00 | -0.23 | 0.82 |

**Table S. 10.** Main effects of the mean beta power and pseudo-*t*-value**.**

| contrast | estimate | SE | df | *t*-value | *p*-value |
|---|---|---|---|---|---|
| $\bar{\boldsymbol{P}}_{\boldsymbol{PMC_L}}(\boldsymbol{\beta})$ younger – older | 3.19 | 1.19 | 33.00 | 2.69 | 0.01 |
| $\bar{\boldsymbol{P}}_{\boldsymbol{M1_L}}(\boldsymbol{\beta})$ normal – difficult (LHDL) | -0.19 | 0.06 | 99.00 | -2.91 | $<10^{-3}$ |
| $\bar{\boldsymbol{P}}_{\boldsymbol{M1_R}}(\boldsymbol{\beta})$ normal – difficult (LHDL) | -0.22 | 0.07 | 99.00 | -3.24 | $<10^{-3}$ |
| $\bar{\boldsymbol{P}}_{\boldsymbol{pMC_L}}(\boldsymbol{\beta})$ normal – difficult (LHDL) | -0.20 | 0.07 | 99.00 | -2.91 | $<10^{-3}$ |
| $\bar{\boldsymbol{P}}_{\boldsymbol{PMC_R}}(\boldsymbol{\beta})$ normal – difficult (LHDL) | -0.23 | 0.07 | 99.00 | -3.03 | $<10^{-3}$ |
| $\bar{\boldsymbol{t}}_{\boldsymbol{M1_L}}(\boldsymbol{\beta})$ normal – difficult (RHDL) | 0.03 | 0.01 | 99.00 | 3.30 | $<10^{-3}$ |
| $\bar{\boldsymbol{t}}_{\boldsymbol{M1_R}}(\boldsymbol{\beta})$ normal – difficult (RHDL) | 0.02 | 0.01 | 99.00 | 2.52 | 0.01 |
| $\bar{\boldsymbol{t}}_{\boldsymbol{pMC_L}}(\boldsymbol{\beta})$ normal – difficult (RHDL) | 0.03 | 0.01 | 99.00 | 3.15 | $<10^{-3}$ |
| $\bar{\boldsymbol{t}}_{\boldsymbol{PMC_R}}(\boldsymbol{\beta})$ normal – difficult (RHDL) | 0.03 | 0.01 | 99.00 | 2.90 | $<10^{-3}$ |

**Table S.11.** Interaction effects of the mean beta power and pseudo-*t*-value.

[†] Abbreviations: Standard error (SE); degree of freedom (df), left hand difficulty level (LHDL); right hand difficulty level (RHDL).

| contrast | LHDL | estimate | SE | df | *t*-value | *p*-value |
|---|---|---|---|---|---|---|
| $\bar{t}_{M1_L}(\beta)$ normal – difficult (RHDL) | normal | 0.07 | 0.01 | 99.00 | 4.52 | $<10^{-3}$ |
| $\bar{t}_{M1_L}(\beta)$ normal – difficult (RHDL) | difficult | 0.00 | 0.01 | 99.00 | 0.15 | 0.88 |
| $\bar{t}_{M1_R}(\beta)$ normal – difficult (RHDL) | normal | 0.05 | 0.01 | 99.00 | 3.54 | $<10^{-3}$ |
| $\bar{t}_{M1_R}(\beta)$ normal – difficult (RHDL) | difficult | 0.00 | 0.01 | 99.00 | 0.02 | 0.98 |
| $\bar{t}_{PMC_L}(\beta)$ normal – difficult (RHDL) | normal | 0.06 | 0.01 | 99.00 | 4.22 | $<10^{-3}$ |
| $\bar{t}_{PMC_L}(\beta)$ normal – difficult (RHDL) | difficult | 0.00 | 0.01 | 99.00 | 0.24 | 0.81 |

### Association between motor behavior and spectral activities

**Table S.12.** LME *t*-test indicating the direction of the significantly associated pseudo-*t*-values with the performance error during unimanual performance.

| | estimate | SE | df | *t*-value | *p*-value |
|---|---|---|---|---|---|
| $\bar{t}_{M1_L}(\beta)$ | -0.271 | 0.360 | 111.918 | -0.752 | 0.45 |
| $\bar{t}_{M1_R}(\beta)$ | -0.405 | 0.509 | 119.937 | -0.795 | 0.43 |
| $\bar{t}_{prec_R}(\alpha)$ | -0.714 | 0.375 | 82.108 | -1.905 | 0.06 |

**Table S.13.** LME *t*-test indicating the direction of the significantly associated pseudo-*t*-values with the performance error during bimanual performance.

| | estimate | SE | df | *t*-value | *p*-value |
|---|---|---|---|---|---|
| $\bar{t}_{M1_L}(\beta)$ | -1.572 | 0.767 | 16.962 | -2.049 | 0.06 |
| $\bar{t}_{M1_R}(\beta)$ | -1.495 | 0.644 | 67.834 | -2.321 | 0.02 |
| $\bar{t}_{PMC_L}(\beta)$ | -2.046 | 1.032 | 28.894 | -1.983 | 0.06 |
| $\bar{t}_{prec_L}(\alpha)$ | -0.948 | 0.591 | 49.367 | -1.605 | 0.12 |
| $\bar{t}_{prec_R}(\alpha)$ | -1.196 | 0.620 | 105.859 | -1.929 | 0.06 |
